# A circuit-to-muscle signaling axis controls locomotor gait transitions in *C. elegans*

**DOI:** 10.64898/2026.04.10.717587

**Authors:** Kyeong Min Moon, Jihye Cho, Jimin Kim, Kyuhyung Kim

**Affiliations:** Department of Brain Sciences, DGIST, Daegu, 42988, Republic of Korea

**Keywords:** *C. elegans*, Gait transition, SMB neuron, NALCN, *unc-79*, nAChR, *unc-29*

## Abstract

Animals switch between locomotor gaits to adapt to changing environments, yet how defined neural circuits and genes coordinate gait transitions remains poorly understood. Here, we show that the head motor neuron SMB acts as a critical gating node that enables the transition from crawling to swimming in *Caenorhabditis elegans*. SMB ablation disrupts head–body synchronization, prolongs muscle activation, and impairs curvature switching during swimming, indicating that the SMB neurons function to constrain motor output to stabilize swimming-specific oscillations. Through an unbiased forward genetic screen, we identify the NALCN channel complex component UNC-79 and the muscle nicotinic acetylcholine receptor subunit UNC-29 as key molecular determinants of gait transition. UNC-79 functions across multiple head interneuron modules to stabilize circuit-wide excitability required for sustained swimming, whereas UNC-29 acts cell-autonomously in body wall muscles to refine calcium amplitude and propagation dynamics. Genetic epistasis analyses position the SMB neurons upstream of UNC-29–mediated muscle activation, establishing a circuit-to-muscle signaling axis that links defined head motor neurons to downstream molecular effectors. Together, our findings show that gait transition emerges from coordinated modulation of neuronal excitability and muscle calcium dynamics, providing a mechanistic framework for how neural circuits reconfigure motor states to adapt locomotor behavior.

**HIGHLIGHTS:**

- The SMB head motor neurons govern the crawl-to-swim transition
- UNC-79/NALCN stabilizes circuit excitability for sustained swimming
- Muscle nAChR subunit UNC-29 tunes calcium gain during gait transition
- A circuit-to-muscle pathway coordinates locomotor state transition

**IN BRIEF:** Gait transitions require coordinated changes in neural circuit activity and muscle output. Moon et al. show that the SMB head motor neurons regulate the switch from crawling to swimming in *C. elegans* by coupling distributed circuit excitability to muscle calcium gain control, defining a circuit-to-muscle pathway that stabilizes locomotor state transitions.

## INTRODUCTION

Animals continuously adjust their locomotion in response to changes in external and internal conditions, allowing them to effectively adapt to ever-changing environments and physiological state ^1–3^. The stereotyped pattern of body movements that drives locomotion is referred to as a gait ^4^. Animals often deploy more than one gait and transition between them to optimize locomotion, yet the strategies governing gait transitions differ markedly across species and even within a single organism ^1–3^. For example, in humans, gait transitions involve shifts between distinct patterns of bipedal locomotion, such as walking and running, typically determined by speed and energetic efficiency ^5–7^. These transitions arise from coordinated interactions between sensory and motor circuits. However, the molecular and circuit mechanisms that enable such gait transitions remain poorly understood.

A multitude of studies has begun to elucidate the mechanisms underlying gait regulation and gait transitions. In mice, locomotion comprises speed-dependent gait types, including alternating gaits such as walking and trotting at lower speeds and synchronous gaits including galloping and bounding at higher speeds ^8,9^. These distinct gait patterns are controlled by discrete populations of glutamatergic neurons in the midbrain. Glutamatergic neurons in both the cuneiform nucleus (CnF) and pedunculopontine nucleus (PPN) regulate low-speed alternating gaits, whereas only CnF glutamatergic neurons initiate high-speed synchronous gaits ^10^, indicating that coordinated activity between CnF and PPN circuits governs gait transitions across locomotor speeds. Like limbed vertebrates, *Drosophila* relies on multi-jointed legs for walking and adjusts its walking speed in response to environmental stimuli ^11,12^. For example, flies increase their speed in response to startling stimuli, a process modulated by serotonergic neurons and receptors within locomotor circuits in the ventral nerve cord ^13^. However, the precise mechanisms by which neuronal activation governs gait and coordinates distinct muscle activity remain poorly understood.

The nematode *Caenorhabditis elegans* is an ideal model system for studying gait transitions, due to its relatively simple nervous system with 302 neurons and a fully mapped synaptic wiring of its motor circuits ^14,15^. Additionally, *C. elegans* exhibits well-defined and adaptable locomotive behaviors. For instance, it displays distinct gaits, crawling and swimming (historically referred to as ‘thrashing’), depending on the physical environment, such as solid surfaces or liquid media. These gaits are characterized by waveforms with distinct amplitudes and wavelengths. On solid agar surfaces, high mechanical friction induces an S-shaped sinusoidal motion with low frequency and short wavelengths ^16^. In contrast, in liquid environments, reduced mechanical friction causes a C-shaped motion with higher frequency and longer wavelengths ^17,18^. During crawling, *C. elegans* undulates at approximately 0.5 Hz, whereas during swimming, it undulates at a significantly higher frequency of 2 Hz ^17–19^. Previous studies have shown that the distinction between crawling and swimming is not solely dictated by mechanical friction but is also modulated by genetic and neurochemical factors. For example, *che-3* mutants, which lack functional ciliated sensory neurons, exhibit defects in transitioning between crawling and swimming ^17^. Additionally, the monoamines serotonin and dopamine play crucial roles in gait switching, such that serotonin promotes the transition from crawling to swimming, whereas dopamine facilitates the switch from swimming to crawling ^18^. However, the precise neural circuit mechanisms and molecular pathways that regulate muscle activity in response to environmental cues during gait transitions remain poorly understood.

Here, we show that the *C. elegans* head motor neuron SMB plays a central role in coordinating gait transitions by regulating muscle activity during swimming. Genetic ablation of the SMB neurons results in prolonged and excessive neck muscle activation, leading to impaired curvature switching and defective crawl-to-swim (C-S) transitions. Through an unbiased forward genetic screen, we identify two molecular determinants of gait transition: the nicotinic acetylcholine receptor (nAChR) subunit UNC-29 and the sodium leak channel complex component UNC-79. UNC-29 acts cell-autonomously in body wall muscles to modulate calcium dynamics required for efficient swimming, whereas UNC-79 functions in head neuronal circuits, including the RMD motor neurons and the the RID interneuron, to regulate locomotor state transitions. Genetic epistasis analyses further support a functional interaction between the SMB motor neurons and UNC-29–mediated muscle activation. At the functional level, the SMB head motor neurons regulate muscle activation through UNC-29 in body wall muscles, while UNC-79 activity in the RMD motor neurons and the RID interneuron modulates body bending to coordinate gait transitions. Together, these findings establish a circuit-to-muscle signaling axis that links defined head motor neurons to downstream molecular effectors and muscle calcium dynamics to ensure precise gait transitions in *C. elegans*.

## RESULTS

### *C. elegans* employs distinct locomotion strategies and muscle activity patterns for crawling and swimming

*C. elegans* exhibits distinct locomotion patterns or gaits depending on the physical environment such that it crawls on soft or solid surfaces and swims in liquid ^17,20,21^ (Figure 1A). Consistent with previous reports ^17,18^, we observed that worms displayed a significantly increased movement rate in liquid, with the swimming frequency being seven times higher than crawling frequency (Figure 1B), indicating that worms may require greater muscle activation for swimming than for crawling. During crawling, worms frequently exhibit head locomotive behaviors, including stomatal oscillation and head lifting ^22^. We found that during swimming, head locomotive behaviors were completely suppressed (Figure 1C), suggesting that swimming utilizes distinct motor patterns.

**Figure 1.**
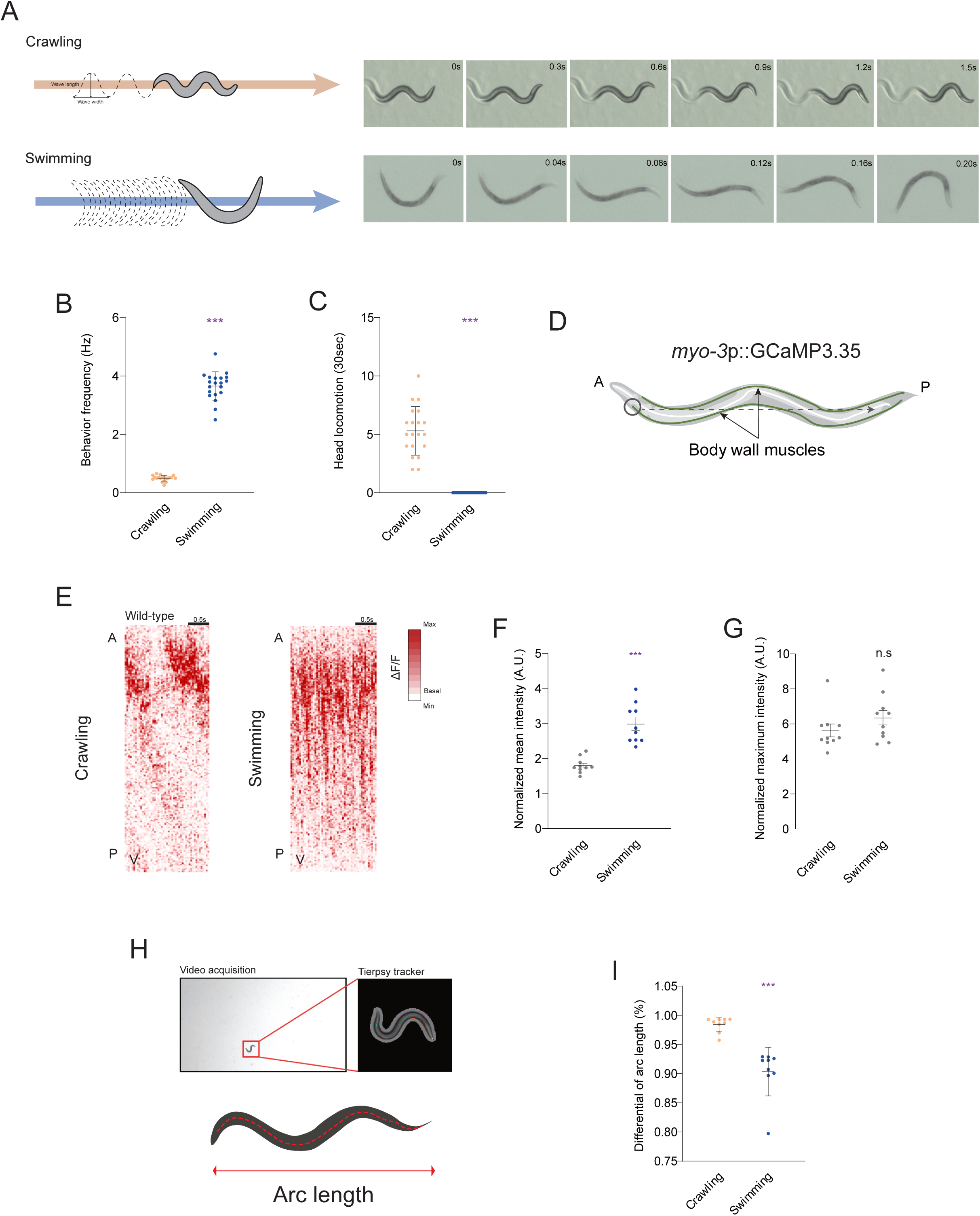
Distinct locomotor modes and muscle activity patterns during crawling and swimming. (A) Schematic illustration and representative time-series images of crawling and swimming behavior. (B) Behavior frequency of wild-type animals during crawling and swimming. n = 20. (C) Number of head locomotion during crawling and swimming. n = 20. (D) Schematic of the experimental setup for monitoring Ca^2+^ activity in body-wall muscles during crawling and swimming. Ca^2+^ signals were recorded from transgenic animals expressing GCaMP3.35 under the control of the *myo-3* promoter (*goeIs3*; see Methods). (E) Representative Ca^2+^ activity heat maps of ventral body-wall muscles in wild-type animals during crawling and swimming. The x-axis denotes time, and the y-axis indicates body position from anterior to posterior. (F) Mean Ca^2+^ fluorescence intensity of body-wall muscles during crawling and swimming. n = 10. (G) Maximum Ca^2+^ fluorescence intensity of body-wall muscles during crawling and swimming. n = 10. (H) Schematic and representative images illustrating worm tracker–based analysis used to measure body arc length. (I) Changes in arc length of wild-type animals during crawling and swimming. n = 9. Data are presented as mean values ± SD (B and C) and ± SEM (F and G). *** indicate a significant difference from wild-type at ***p < 0.001 is shown (unpaired t-test followed by two-tailed test).

The body wall muscles of *C. elegans* form a four-quadrant structure, consisting of two dorsal and two ventral rows, and run continuously along the dorsal and ventral sides in a longitudinal arrangement ^23^ (Figure 1D). We then examined body wall muscle activity patterns in freely moving transgenic animals expressing GCaMP3.35 in all body wall muscles ^24^ (Figure 1D). Consistent with previous observations, we observed that during crawling, Ca^2+^ activity in the body wall muscles is strongly localized to specific, mainly head, muscle regions and exhibits a wave-like propagation along the body from the head to the tail (Figure 1E). In contrast, Ca^2+^ activity during swimming appears simultaneously across the entire body wall muscles without a distinct propagation pattern (Figure 1E). This suggests that crawling relies on sequentially propagating muscle activation, whereas swimming involves increased muscle activity throughout the whole-body wall muscles. We also found that while the maximum Ca²⁺ intensity did not differ between crawling and swimming, the overall Ca^2+^ signals in the body wall muscles were significantly higher during swimming (Figures 1F–1G), confirming increased muscle activity in swimming. We also measured the body length of worms along the central body axis during both crawling and swimming (Figure 1H) and found that body length was reduced during swimming compared to crawling (Figure 1I), suggesting that *C. elegans* employs greater muscle contraction to bend its body and generate the C-shaped posture characteristic of swimming behavior.

### The SMB and SMD head motor neurons mediate swimming by regulating posture transitions

The SMB and SMD neurons consist of two pairs of neurons with cell bodies located in the head, with their processes extending sublaterally from the head to the tail to innervate head muscles ^15^. These neurons have been shown to mediate sinusoidal undulatory locomotion in *C. elegans* ^25,26^ (Figures 2A and S1A). To determine whether these head motor neurons mediate crawling as well as swimming, we genetically ablated the SMB and SMD neurons by expressing *Caspase* genes under the control of cell-specific promoters ^27^ and quantified crawling and swimming frequency. We found that both SMB- and SMD-ablated worms exhibited decreased crawling and swimming frequencies (Figures 2B–2C). Moreover, in *lim-4* mutants, where SMB neurons are not properly specified and thus nonfunctional ^26,28,29^, we observed similar defects in crawling and swimming behavior as in SMB-ablated worms (Figures S1B–S1C). However, ablation of the AWB neurons, which are also disrupted in *lim-4* mutants, did not result in defects in crawling or swimming (Figures S1D–S1E). These data indicate that the SMB and SMD head motor neurons are required for both crawling and swimming in *C. elegans*.

**Figure 2.**
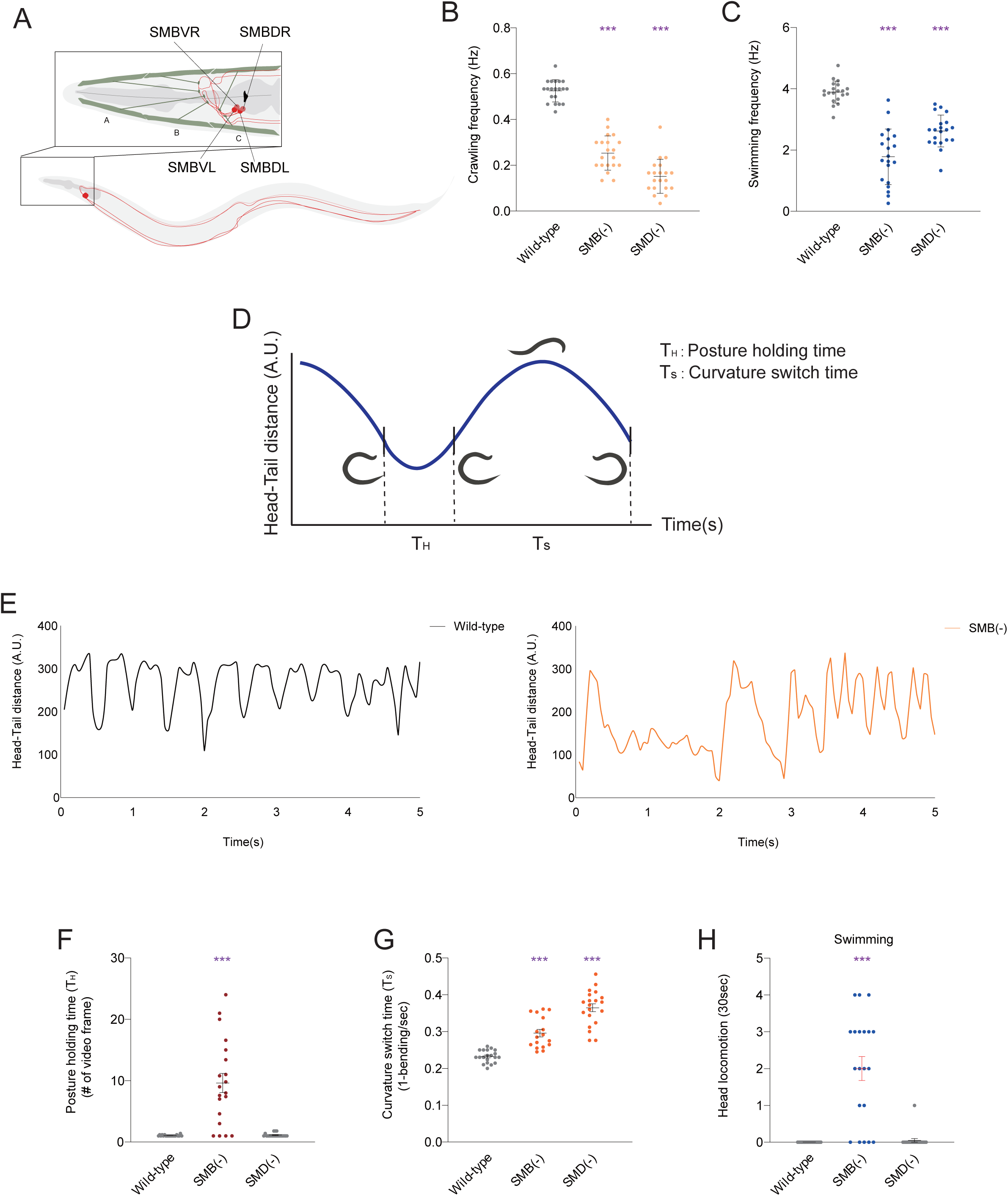
The SMB and SMD head motor neurons differentially control swimming dynamics. (A) Schematic illustrating the SMB head motor neurons in *C. elegans*. (B) Crawling frequency of wild-type animals and animals with genetically ablated SMB or SMD neurons. The SMB neurons were ablated by expressing split-caspase constructs (TU#813 and TU#814) under the control of the *flp-12*Δ3 promoter (KHK340), and The SMD neurons were ablated using the *flp-22* and *lad-2* promoters (KHK1174). n = 20. (C) Swimming frequency of wild-type, SMB (-) and SMD (-). n = 20. (D) Schematic defining curvature switch time and posture holding time during swimming. (E, F) Posture holding time of wild-type, SMB (-) and SMD (-) animals. n = 20. (G) Curvature switch time of wild-type, SMB (-) and SMD (-) animals. n ≥ 17 for each. (H) Total number of head locomotion during swimming in wild-type, SMB (-) and SMD (-) animals. n = 20. Data are presented as mean values ± SD (B, C) and SEM (F, G, and H). *** indicate a significant difference from wild-type at ***p < 0.001 is shown (one-way ANOVA test followed by Dunnett’s multiple comparisons test).

To further characterize factors responsible for the reduced locomotion frequencies observed in SMB- or SMD-ablated worms, we measured two kinematic parameters during swimming, namely curvature switch time (the time taken to transition from one C-shaped body bend to the opposite curvature) and posture holding time (the duration spent in the fully bent C-shape before transitioning) (Figure 2D). SMB-ablated worms exhibited a modest increase in curvature switch time, whereas posture holding time was markedly increased compared to wild-type animals (Figures 2E–2G). In contrast, SMD-ablated worms exhibited an increase in curvature switch time, while posture holding time remained comparable to wild-type levels (Figures 2F–2G, Supplementary Video 1-3). These findings suggest that the SMB neurons play a main role in initiating curvature transitions and, together with the SMD neurons, facilitate postural switching during swimming. We also observed that in SMB-ablated worms, but not in SMD-ablated worms, head locomotion was not suppressed during swimming (Figure 2H), suggesting that the SMB neurons further coordinate the suppression of head-driven motor output during swimming.

Next, we optogenetically stimulated the SMB and SMD neurons by expressing red-shifted channelrhodopsin (ReaChR) ^30,31^ under the control of the *flp-12Δ3* and *flp-22* promoters, which drive expression in the SMB and SMD neurons, respectively (Figure S1F). Chronic activation of either all four SMB or SMD neurons did not alter crawling behavior (Figure S1G). In addition, activation of the SMB neurons, but not the SMD neurons, led to a slight reduction in swimming frequency (Figure S1H). These findings suggest that our optogenetic stimulation conditions may not fully recapitulate the endogenous activity patterns of the SMB and SMD neurons, or that sustained activation of these neurons alone is insufficient to drive behavioral changes.

### The SMB neurons prevent excessive muscle activation to ensure smooth transitions of posture and gait

To investigate the underlying mechanisms by which the SMB and SMD neurons regulate posture transitions, we performed whole-body Ca^2+^ imaging of body wall muscles in freely moving animals during crawling and swimming (Figure 1D). SMB-ablated worms exhibited a significant increase in both the mean and maximum Ca^2+^ activity across the entire body wall muscles compared to controls during both crawling and swimming (Figures 3A–3F and S2A). In contrast, SMD-ablated worms showed no significant changes in both the mean and maximum Ca^2+^ activity, despite similarly reduced locomotion frequency (Figures S2B–S2E). These results suggest that the swimming defects observed in SMB-ablated animals may arise from excessive muscle activation, leading to impaired coordination of posture transitions.

**Figure 3.**
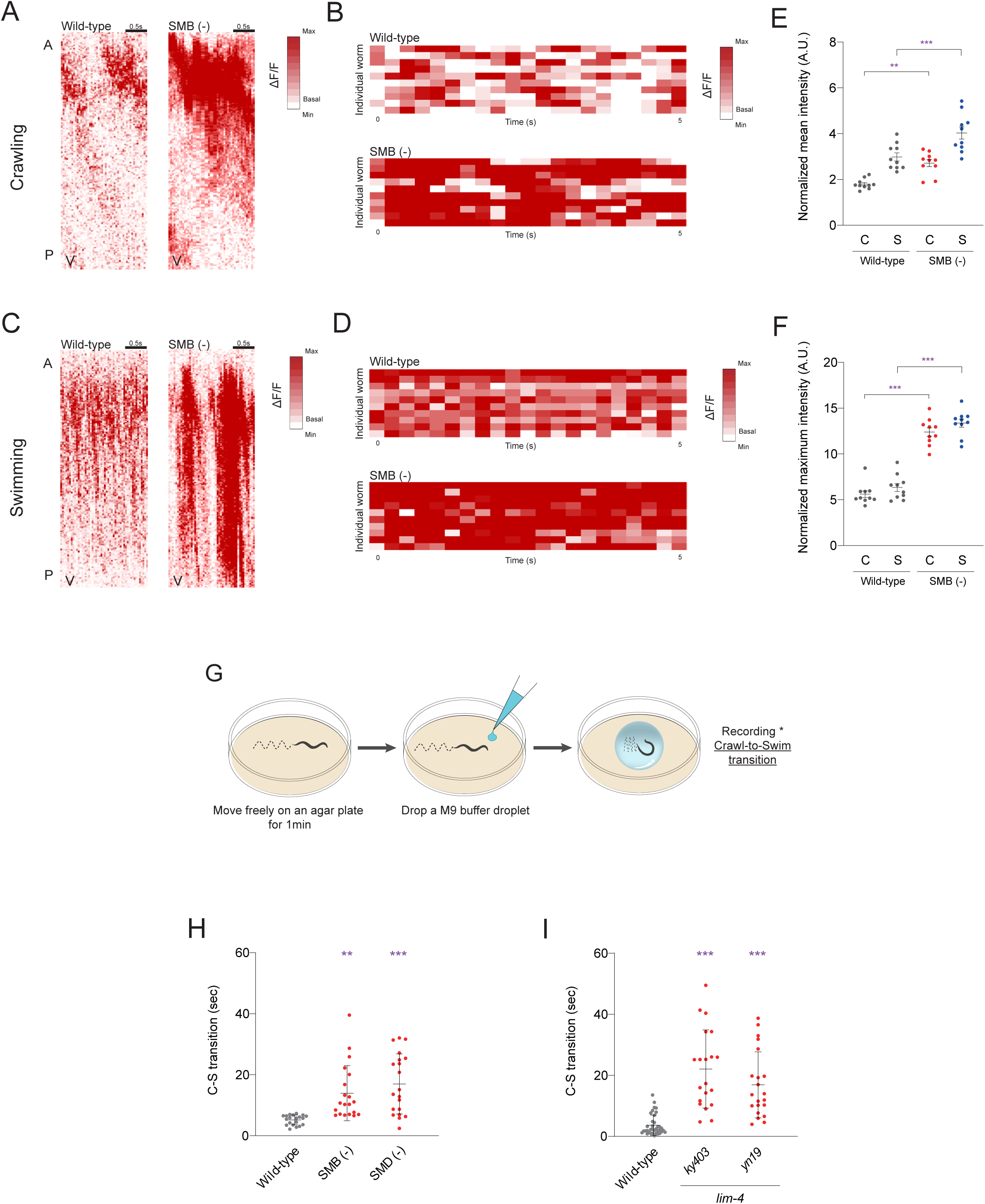
The SMB neurons constrain muscle activity and promote gait transition. (A, C) Representative Ca²⁺ activity heat maps of ventral body-wall muscles in wild-type and SMB (-) animals during crawling and swimming, respectively. The x-axis denotes time, and the y-axis indicates body position from anterior to posterior. (B, D) Heat maps showing ΔF/F of Ca^2+^ activity in neck muscle of wild-type and SMB (-) during crawling and swimming, respectively. n = 10 for each. (E, F) Mean and maximum of Ca^2+^ fluorescence intensity of body-wall muscles in wild-type and SMB (-) animals during crawling and swimming (C denotes crawling; S denotes swimming). n = 10 for each. (G) Schematic of the crawl-to-swim transition assay. (H) Crawl-to-swim transition time of wild-type, SMB (-), and SMD (-) animals. n = 20. (I) Crawl-to-swim transition time of wild-type, *lim-4* mutant alleles (*ky403*, *yn19*). n ≥ 20. Data are represented ± SEM. ** and *** indicate a significant difference from wild-type at **p < 0.01 and ***p < 0.001 are shown (one-way ANOVA test followed by Dunnett’s multiple comparisons test).

Given the distinct effects of SMB and SMD ablation on muscle activation, we next asked whether these neurons are also required for coordinating gait transitions between crawling and swimming and performed a crawl-to-swim (C–S) transition assay (see Methods for details; Figure 3G). We found that worms with ablated SMB or SMD neurons exhibited a significant delay in C-S transition times (Figure 3H). To further confirm the role of the SMB neurons in gait transitions, we performed the C-S assay in two *lim-4* mutant alleles (*ky403* and *yn19*), as well as in worms with genetically ablated AWB neurons. Similar to SMB-ablated worms, *lim-4* mutants displayed impaired gait transitions (Figure 3I), whereas AWB-ablated animals showed no such defects (Figure S3A). These results indicate that the SMB and SMD neurons facilitate the transition from crawling to swimming in *C. elegans* by sustaining locomotion frequencies required for efficient gait switching.

The SMB neurons have been shown to regulate head locomotion through the neuropeptide FLP-12 ^22^. We next asked whether the SMB neurons might also use FLP-12 to control posture and gait transitions during swimming behavior. We found that although *flp-12* mutants exhibited impaired head locomotion, they showed normal crawl-to-swim transitions and swimming behavior ^22^ (Figure S3B). These results indicate that the SMB neurons regulate gait transition and swimming behavior through mechanisms independent of *flp-12*, possibly involving other neurotransmitters such as acetylcholine ^32^.

### A genetic screen identifies *unc-79* and *unc-29* as molecular determinants of crawl-to-swim transition

The neural mechanisms involving the SMB and SMD neurons suggest a circuit-level control of gait transition, but the molecular factors underlying this regulation remain unknown. Because the ability to switch between crawling and swimming provides a direct behavioral readout of gait regulation, we performed an unbiased genetic screen to isolate mutants defective in crawl-to-swim transitions (Figure 4A). In this screen, we excluded animals with severe crawling defects to isolate mutants specifically impaired in gait transition rather than general locomotor ability. From this screen, we identified three mutant alleles (named *lsk57*, *lsk59*, and *lsk67*) that exhibited delayed crawl-to-swim transitions (Figure 4B). We then further analyzed their crawling and swimming behaviors and found that although we tried to exclude mutants with overt crawling defects, all three alleles nonetheless showed significantly reduced locomotion frequencies compared to wild-type animals (Figures 4C–4D).

**Figure 4.**
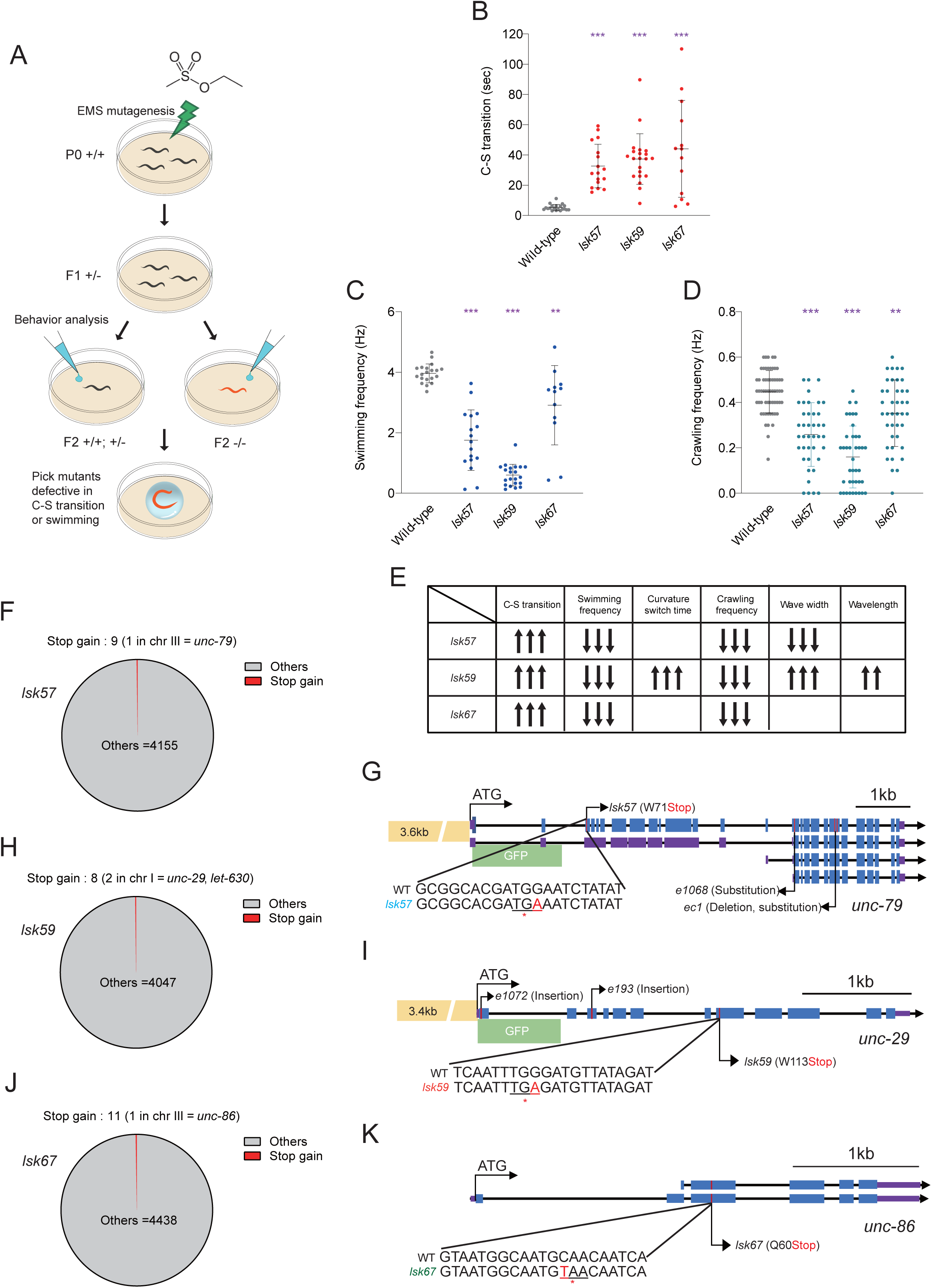
Forward genetic screen identifies regulators of gait transition. (A) Schematic of the EMS mutagenesis–based forward genetic screen. A crawl-to-swim (C–S) transition assay was used to isolate mutant alleles with defects in crawling behavior, gait transition and swimming behavior. (B, C) Crawl-to-swim transition time and swimming frequency of wild-type animals and isolated mutants (*lsk57*, *lsk59*, *lsk67*). n ≥ 12 for each. (D) Crawling frequency of wild-type animals and isolated mutants. n = 40 for each. (E) Summary table of behavioral phenotypes observed in the isolated mutant strains. Black arrows indicate significant defects compare with wild-type. (F, H, J) Pie charts showing the distribution of stop-gain mutations across the coding regions of *lsk57*, *lsk59*, and *lsk67*, respectively. (G, I, K) Genomic structures and promoter regions of *unc-79*, *unc-29*, and *unc-86*. Blue (or purple) boxes and black lines represent exons and introns, respectively. Yellow boxes indicate promoter regions fused to the green fluorescence protein reporter gene. Red lines denote the positions of point mutations identified in the *lsk57*, *lsk59*, *lsk67* respectively. Data are represented ± SD (B, C, D). ** and *** indicate a significant difference from wild-type at **p < 0.01 and ***p < 0.001 are shown (one-way ANOVA test followed by Dunnett’s multiple comparisons test).

Next, we examined body posture during both crawling and swimming. We first quantified body wavelength and wave width during crawling (Figure 1A) and found that *lsk67* mutants were indistinguishable from wild-type animals, whereas the remaining mutant alleles exhibited significant differences in wavelength, wave width, or both compared to wild-type animals (Figures S4A–S4B). Notably, locomotion defects in *lsk57* mutants became progressively more severe over successive generations, eventually resulting in an uncoordinated (Unc) phenotype (Figure S4C). During swimming, we quantified the angle formed by the head, vulva, and tail positions as a measure of body curvature across a single swimming cycle (Figure S5A), and found that the other alleles were comparable to wild-type animals, whereas *lsk57* mutants exhibited severe defects in swimming posture, characterized by a strong bias toward ventral bending, with markedly reduced dorsal bending during the swimming cycle (Figure S5B). Together, these data suggest that defective crawl-to-swim transitions arise from allele-specific disruptions in locomotor kinematics and posture (Figure 4E).

To molecularly define the genetic lesions underlying the observed gait-transition phenotypes, we next subjected the isolated mutants to positional cloning and genome-wide analysis. Initial chromosomal mapping placed *lsk57*, *lsk59*, and *lsk67* on chromosomes III, I, and III, respectively (Figures S6A–S6C). We then performed whole-genome sequencing of each mutant strain (average coverage ∼50×; BGI Genomics), which identified more than 4,000 single-nucleotide polymorphisms (SNPs per genome). Notably, each mutant harbored one or two stop-gain mutations within the genetically mapped chromosome (Figures 4F, 4H, and 4J). We then identified the causative molecular lesions in *lsk57*, *lsk59*, and *lsk67* as premature stop codons in *unc-79* (W71Stop), *unc-29* (W113Stop), and *unc-86* (Q60Stop), respectively (Figures 4G, 4I, 4K and S7A–S7C). We decided to further characterize *unc-79* and *unc-29* but not the *unc-86* gene, as mutations in *unc-86* are known to cause broad developmental defects in the nervous system, complicating interpretation of its role in locomotor gait regulation ^33,34^.

### The NALCN channel complex, including UNC-79, is required for efficient gait transition and swimming behavior

The *unc-79* gene encodes an auxiliary subunit of the NALCN sodium leak channel complex, which forms a functional protein complex with UNC-80 and is required for proper expression and axonal localization of the NALCN channels NCA-1 and NCA-2 ^35–37^ (Figure 5A). In *C. elegans*, NALCN channels function in establishing neural resting potential that stabilizes neuronal excitability and sustains locomotor activity ^35,36^.

**Figure 5.**
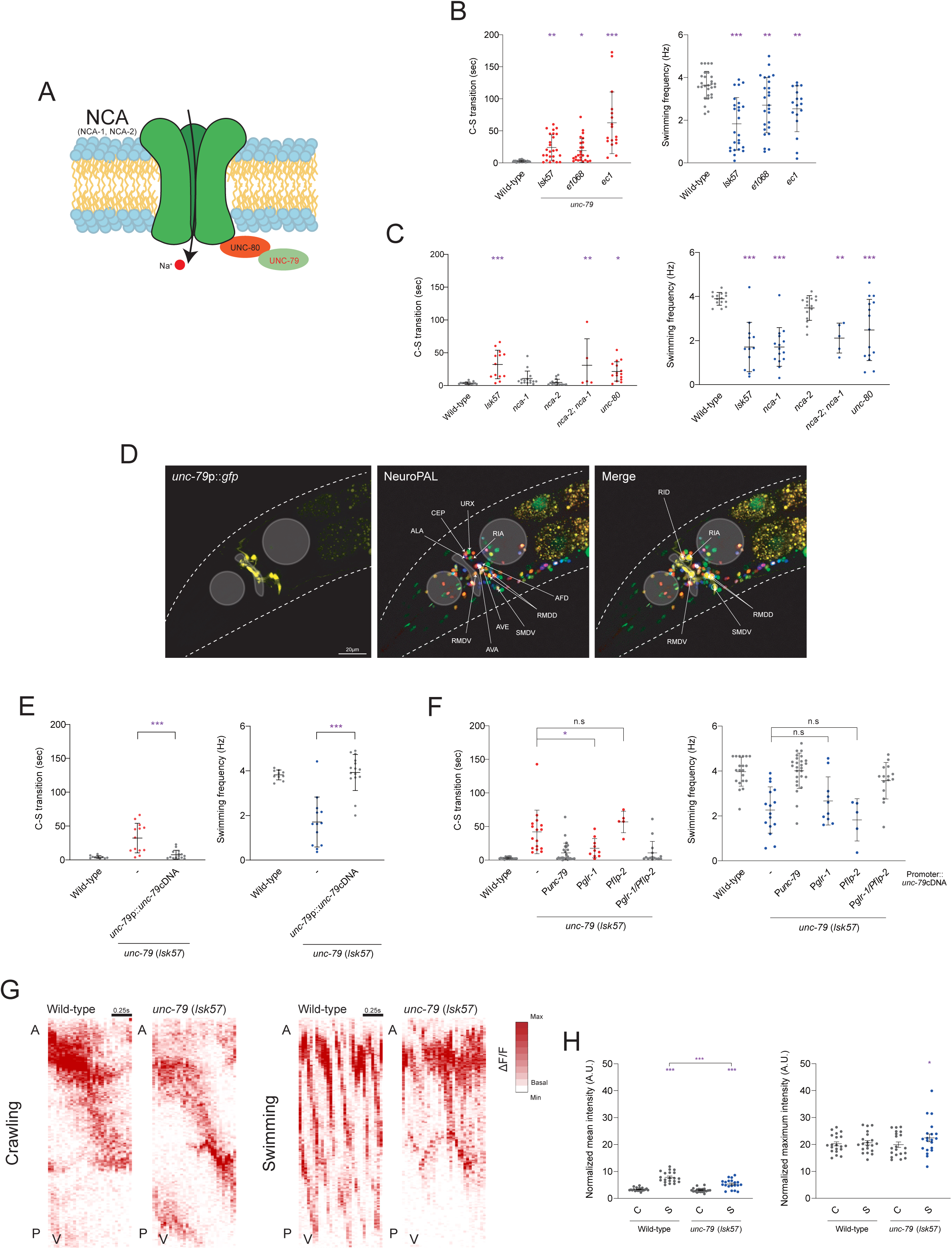
UNC-79/NALCN signaling in head interneurons regulates gait transition and swimming. (A) Schematic representation of the NCA channel complex in *C. elegans*. (B) Crawl-to-swim (C–S) transition time and swimming frequency of *unc-79* mutant alleles (*lsk57*, *e1068*, *ec1*). n ≥ 17 for each. (C) Crawl-to-swim transition time and swimming frequency of wild-type animals and NCA channel defective mutants (*nca-1*, *nca-2*, and *unc-80*) (n ≥ 13 animals per genotype, except nca-1 and *nca-2*, which exhibit severe locomotor defects; n = 5). (D) Neuronal cell identification of *unc-79*-expressing neurons using NeuroPAL worm. The merge images of transgenic animals co-expressing NeuroPAL and *unc-79*p::*gfp* (yellow) in the head. (E) Crawl-to-swim transition time and swimming frequency of wild-type and *lsk57* mutant animals expressing *unc-79* cDNA under the control of the *unc-79* long-isoform promoter. n ≥ 10 for each. (F) Crawl-to-swim transition time and swimming frequency of wild-type and *lsk57* mutant animals expressing *unc-79* cDNA under the control of interneuron-specific promoters (*glr-1* and *flp-2*). n ≥ 5 for each. (G) Representative Ca^2+^ activity heat maps of ventral body-wall muscles in wild-type and *unc-79* mutant animals during crawling and swimming. The x-axis denotes time, and the y-axis indicates body position from anterior to posterior. (H) Mean and maximum of Ca^2+^ fluorescence intensity of body-wall muscles in wild-type and *lsk57* animals during crawling and swimming, respectively. n = 20 for each. Data are represented ± SD (B–F), SEM (H). *, ** and *** indicate a significant difference from wild-type at *p < 0.05, **p < 0.01 and ***p < 0.001 are shown (one-way ANOVA test followed by Dunnett’s multiple comparisons test (B, C, H) and Tukey’s multiple comparisons test (E, F)).

To validate *unc-79* as the gene underlying the locomotor defects identified in our genetic screen, we examined two independent loss-of-function alleles, *unc-79* (*e1068*) and *unc-79* (*ec1*) (PMID: 2900611, PMID: 3576211) (Figure 4G). Consistent with the phenotype of the *lsk57* mutant allele, both *unc-79* mutant alleles exhibited a significant delay in crawl-to-swim (C–S) transition time and a reduced swimming frequency (Figure 5B), indicating that *unc-79* mediates efficient gait transition and swimming behavior.

We next asked whether other components of the NALCN channel complex exhibit phenotypes similar to those of *unc-79* mutants. To this end, we examined mutants of the functionally redundant NALCN channel genes *nca-1* and *nca-2*, including both single and double mutants, as well as *unc-80* mutants ^37,38^. We found that *nca-2; nca-1* double mutants and *unc-80* mutants exhibited a pronounced delay in crawl-to-swim (C–S) transition and a significant reduction in swimming frequency, resembling the phenotypes observed in *unc-79* mutants (Figure 5C). Notably, *nca-1* or *nca-2* single mutants showed no detectable defects in C–S transition, with only minor effects on swimming frequency observed in *nca-1* mutants (Figure 5C). These results indicate that components of the NALCN channel complex act together to regulate gait transitions and swimming behavior in *C. elegans*.

### UNC-79 acts in multiple neuronal classes in the head to regulate gait transition and swimming-specific muscle activation

To determine the site of *unc-79* action, we generated transgenic animals expressing *unc-79*p::*gfp* under the control of a 3,665-bp upstream promoter region (Figure 4G). Consistent with previous reports (Humphrey et al., 2007), *unc-79* expression was broadly detected in head neurons (Figure 5D). To precisely identify *unc-79*–expressing neurons further, we compared GFP expression with NeuroPAL reference signals ^39^ and observed consistent expression in several neurons of the head, including the neck motor neurons RMD, the RIA and RID interneurons, and the SMD motor neurons (Figure 5D). Expression in RMD, SMD, and RID neurons was further confirmed by colocalization with *glr-1*p::*mCherry* which labels a subset of interneurons and motor neurons including these cell types ^40,41^ (Figure S8A).

To test whether *unc-79* functions within these neurons to regulate gait transition and swimming, we expressed *unc-79* cDNA under the control of the endogenous *unc-79* promoter in *unc-79* mutant animals. This transgene fully rescued crawl-to-swim transition time, swimming frequency, and crawling defects (Figures 5E and S8B–S8D), indicating that *unc-79* acts cell-autonomously. We next performed cell-type–specific rescue experiments by expressing *unc-79* cDNA under the control of the *glr-1* promoter, which is active in RMD, SMD, and several command interneurons, or the *flp-2* promoter, which is expressed in RID ^42^. Expression driven by either promoter alone was insufficient to rescue the *unc-79* mutant phenotypes (Figure 5F). In contrast, combined expression using both *glr-1* and *flp-2* promoters fully restored gait transition and swimming behavior (Figure 5F). Together, these results indicate that *unc-79* acts in multiple neuronal classes, including the RMD/SMD motor neurons and the RID interneuron, to regulate gait transition and swimming behavior.

We next examined body wall muscle activity in *unc-79* mutants during locomotion. During crawling, *unc-79* mutants exhibited largely normal calcium activity in body wall muscles, as indicated by comparable spatiotemporal heat map patterns and mean Ca²⁺ activity levels compared to wild-type animals (Figures 5G–5H). In contrast, during swimming, *unc-79* mutants displayed markedly disrupted muscle activation patterns accompanied by a significant reduction in mean Ca²⁺ activity (Figures 5G–5H). These results indicate that *unc-79* is dispensable for muscle recruitment during crawling but is required for proper activation of body wall muscles during swimming.

### The nAChR subunit UNC-29 mediates efficient gait transition and swimming behavior

The *unc-29* gene encodes a non-alpha subunit of the nicotinic acetylcholine receptor (nAChR) ^43,44^ (Figures 4I and 6A). To further characterize the functional role of *unc-29*, we examined two additional loss-of-function alleles, *unc-29* (*e193*) and *unc-29* (*e1072*) (Figure 4I). Consistent with the *lsk59* mutant, both alleles exhibited a significant delay in the crawl-to-swim transition time and a marked reduction in swimming frequency (Figure 6B). To confirm that the observed locomotor defects were indeed caused by the disruption of *unc-29*, we performed a rescue test by expressing a genomic fragment of *unc-29* and found that the genomic transgene successfully restored both the gait transition delay and the swimming defects in *lsk59* mutants (Figure 6C). These results indicate that *unc-29* is required for efficient gait transitions and sustained swimming activity.

**Figure 6.**
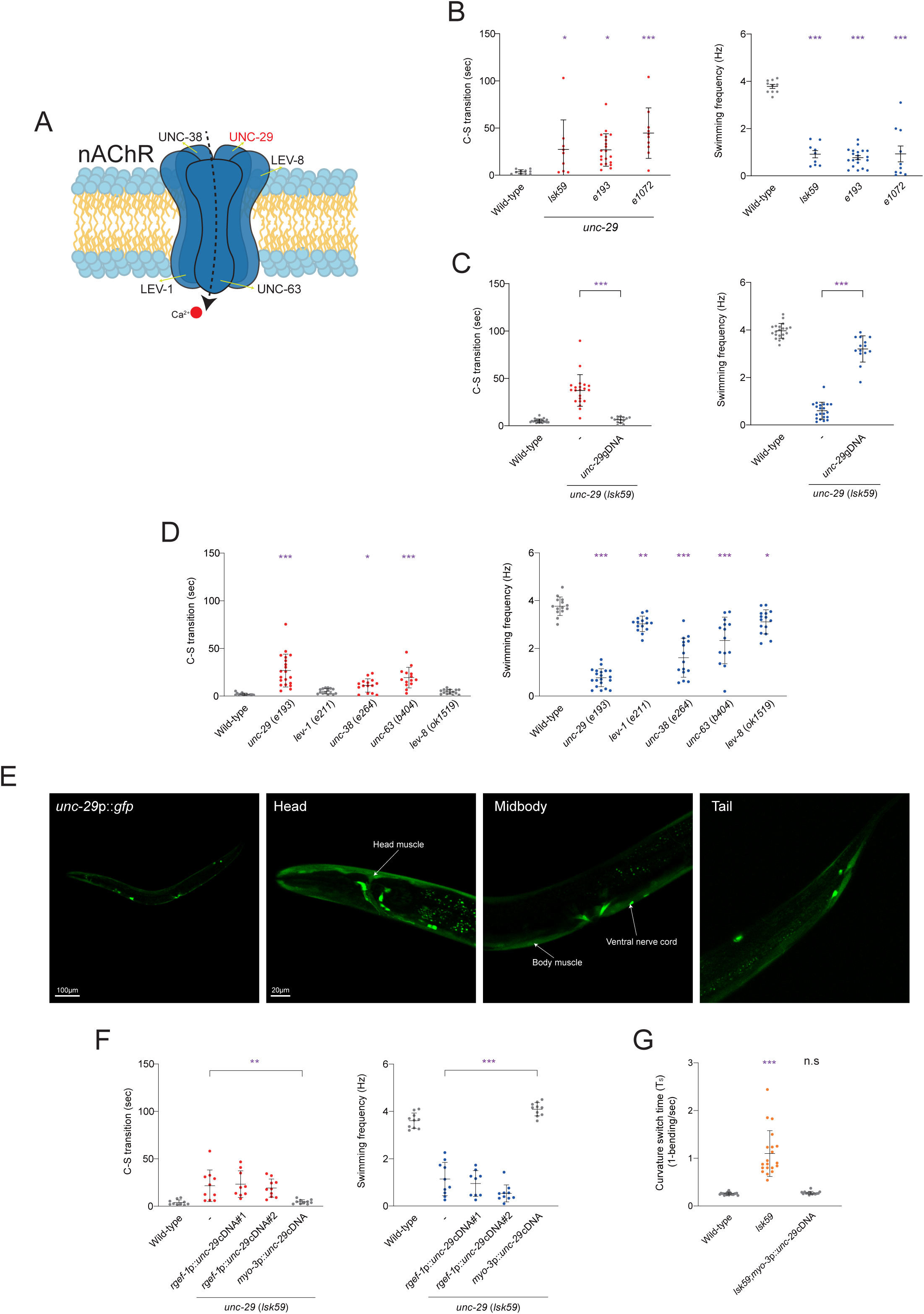
Muscle nAChR subunit UNC-29 regulates gait transition and swimming. (A) Schematic representation of the nAChR channel complex in *C. elegans*. (B) C-S transition time and swimming frequency of wild-type, *lsk59,* and *unc-29* mutant alleles (e*193*, e*1072*). n ≥ 9 for each. (C) Crawl-to-swim transition time and swimming frequency of wild-type and *lsk59* mutant animals expressing *unc-29* gDNA. n ≥ 14 for each. (D) C-S onset time and swimming frequency of wild-type, *unc-29* (*e193*), and nAChR channel subunit mutants (*lev-1* (*e211*), *unc-38* (*e264*), *unc-63* (*b404*), and *lev-8* (*ok1519*)). n ≥ 14 for each. (E) Representative image of a wild-type animal expressing *unc-29*p::*gfp*. *unc-29* is expressed in head and tail neurons (head, tail images), ventral nerve cord (body image), and body muscles (head, body, tail images). Scale bar = 100μm (20x), 20μm (40x). (F) C-S transition time and swimming frequency of wild-type and *lsk59* expressing *unc-29*cDNA under the control of a pan-neuronal expressed *rgef-1* gene promoter and body wall muscle expressed *myo-3* gene promoter. n ≥ 9 for each. (G) Curvature switch time of wild-type, *lsk59* and *lsk59* rescue worms during swimming. n = 20 for each. Data are represented ± SD. *, ** and *** indicate a significant difference from wild-type at *p < 0.05, **p < 0.01 and ***p < 0.001 are shown (one-way ANOVA test followed by Dunnett’s multiple comparisons test (B, D) and Tukey’s multiple comparisons test (C, G, F)).

In *C. elegans*, the muscle nAChR population primarily consists of the levamisole-sensitive receptor (L-AChR), a heteropentameric complex composed of five subunits: UNC-38, UNC-63, LEV-8, UNC-29, and LEV-1 ^43,44^. To determine whether the other subunits of the L-AChR complex also mediate gait transition, we analyzed mutants for each subunit. Consistent with the crawling defects previously reported (PMID: 9221782), we found that *unc-63* and *unc-38* mutants exhibited significant impairments in both gait transition and swimming frequency (Figure 6D). In contrast, *lev-1* and *lev-8* mutants appeared largely normal under the same conditions (Figure 6D). Together, these results indicate that nAChRs contribute to gait transition and swimming behavior in *C. elegans*.

### UNC-29 acts cell-autonomously in body wall muscles to regulate gait transition by modulating muscle calcium dynamics

Previous studies have reported that *unc-29* is expressed in body wall muscles, the ventral nerve cord, and a subset of head neurons ^45^. Consistent with these findings, transgenic animals expressing *unc-29*p::*gfp* driven by a 3,423-bp promoter fragment exhibited consistent GFP expression in body wall muscles, the ventral nerve cord, and a subset of neurons (Figures 4I and 6E). To identify the site of *unc-29* action in regulating gait transition, we performed cell-specific rescue experiments. We expressed *unc-29* cDNA under either the pan-neuronal promoter (*rgef-1*p) or the body wall muscle-specific promoter (*myo-3*p) ^46–48^. While pan-neuronal expression failed to rescue defects of *unc-29* mutants, muscle-specific expression fully rescued the crawl-to-swim transition delay, swimming frequency, and crawling defects (Figures 6F and S9A–S9B). Together, these results indicate that *unc-29* acts cell-autonomously in body wall muscles to regulate gait transition and swimming behavior. Notably, *unc-29* mutants exhibited a prolonged curvature switch time, consistent with disrupted muscle calcium dynamics, and this phenotype was fully rescued by muscle-specific expression of *unc-29* (Figure 6G).

To investigate how *unc-29* regulates muscle activity during locomotion, we monitored *in vivo* calcium dynamics using a transgenic animal that expresses muscle-specific GCaMP3.35. In *unc-29* mutants, the spatiotemporal propagation of calcium signals was significantly slower than in wild-type animals during both crawling and swimming (Figures 7A–7B). Interestingly, despite the sluggish propagation and locomotor defects, *unc-29* mutants exhibited significantly higher mean Ca^2+^ levels in body wall muscles compared to wild-type animals (Figure 7C). This aberrant calcium accumulation was fully rescued by the muscle-specific expression of *unc-29* cDNA, restoring both the signal propagation speed and mean Ca^2+^ levels (Figure 7C).

**Figure 7.**
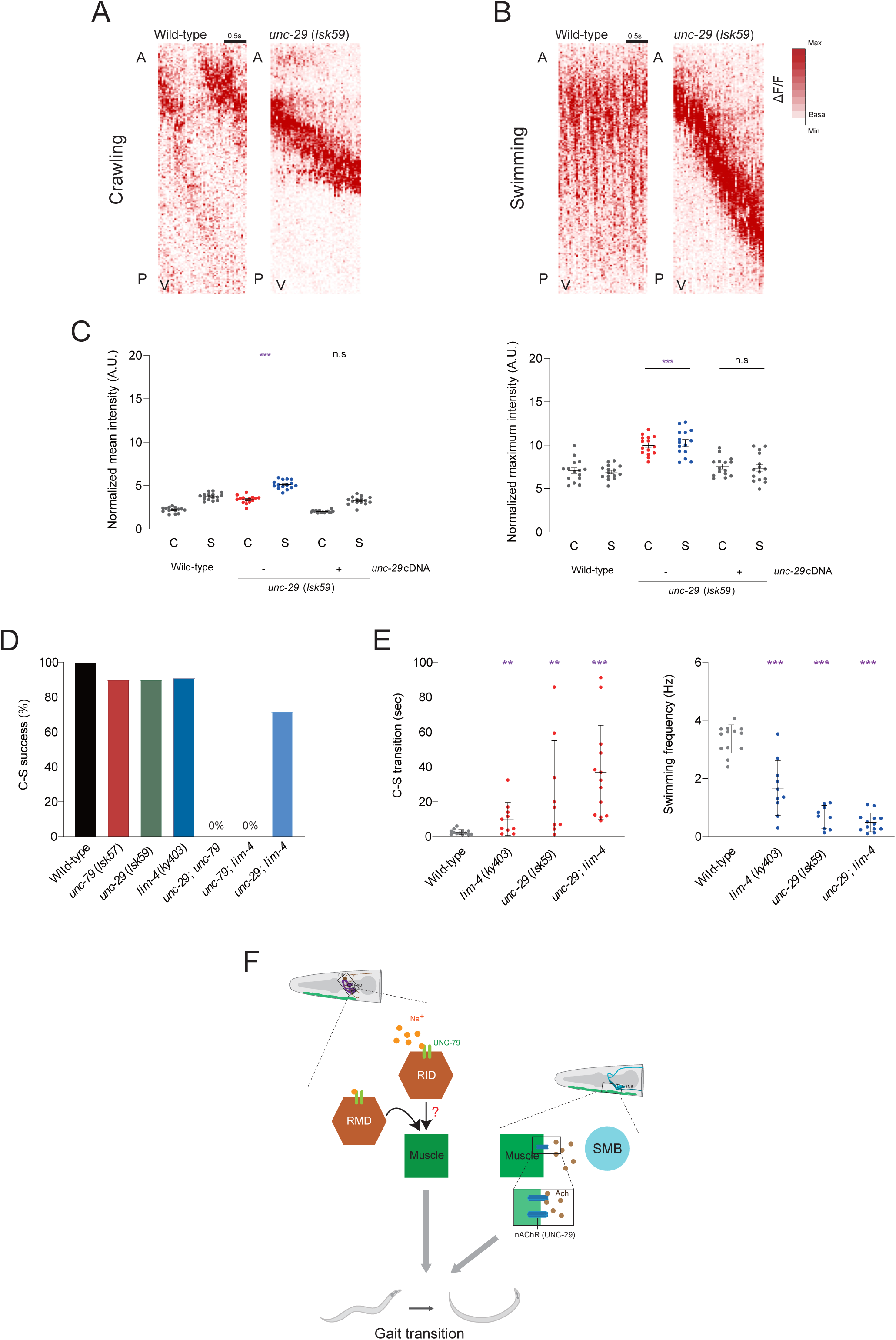
UNC-29 functions downstream of SMB-mediated muscle control during gait transition. (A, B) Representative Ca^2+^ activity heat maps of ventral body-wall muscles in wild-type and *unc-29* mutant animals during crawling and swimming. In (A, B), the x-axis denotes time and the y-axis indicates body position from anterior to posterior. (C) Mean and maximum of Ca^2+^ fluorescence intensity of body-wall muscles in wild-type, *unc-29* mutant, and *unc-29* cDNA expressing animals during crawling and swimming. n = 15 for each. (D) Crawl-to-swim (C–S) transition success rate of wild-type, *lim-4* (*ky403*), *unc-79* (*lsk57*), *unc-29* (*lsk59*), *unc-29*; *unc-79*, *unc-79*; *lim-4*, and *unc-29*; *lim-4* animals. n ≥ 7 for each. (E) Crawl-to-swim transition time and swimming frequency of wild-type, *lim-4*, *unc-29*, and *unc-29*; *lim-4* animals. n ≥ 9 for each. (F) Model illustrating the circuit mechanisms underlying gait transitions in *C. elegans*. Upon encountering liquid, animals initiate gait transitions and sustain swimming behavior via two parallel pathways: an interneuron hub expressing the NALCN channel subunit UNC-79 (including RID and RMD) and the head motor neuron SMB. Swimming is associated with Ca^2+^ activity patterns distinct from those observed during crawling, and the *unc-29* encoded nicotinic acetylcholine receptor may contribute to shaping these muscle activity patterns downstream of the SMB neurons. Data are represented ± SD (E) and SEM (C). ** and *** indicate a significant difference from wild-type at **p < 0.01 and ***p < 0.001 are shown (one-way ANOVA test followed by Tukey’s multiple comparisons test).

### Genetic epistasis analysis suggests that the SMB motor neurons regulate gait transition through UNC-29-mediated muscle activation

To delineate the functional hierarchy between the SMB head motor neurons, UNC-79-expressing interneurons, and UNC-29-containing muscle nAChRs, we performed a genetic epistasis analysis. We first generated three sets of double mutants including *unc-29; unc-79*, *unc-79; lim-4*, and *unc-29; lim-4* and evaluated their locomotor performance using the C–S assay. We observed that both *unc-29; unc-79* and *unc-79; lim-4* double mutants exhibited a synergistic and severe deficit, completely failing to initiate swimming behavior (Figure 7D). This total loss of gait switching capability suggests a potent functional interaction between these components. In contrast, the *unc-29; lim-4* double mutant was still capable of initiating the gait transition but it exhibited a delayed transition time and reduced swimming frequency similar to that of either the *unc-29* or *lim-4* single mutants (Figures 7D–7E). This data suggests that the SMB motor neurons regulate head muscle activity during gait transitions via *unc-29*-expressed nAChRs.

## Discussion

Swimming in *C. elegans* is characterized by a C-shaped body posture with increased frequency and wavelength compared to crawling ^17,19^. One feature we uncovered is that swimming is accompanied by a suppression of independent head exploratory behaviors, resulting in tightly synchronized head–body undulation. This observation suggests that gait transition is not merely a mechanical consequence of reduced friction, but involves an active neural mechanism that reconfigures head motor output to match a swimming-specific motor state. The SMB neurons have previously been implicated in head posture control and exploratory head scanning. Our data extends this role by revealing that the SMB neurons function as a gating module that constrains head motor output during gait switching. Genetic ablation of the SMB neurons not only delayed crawl-to-swim transitions but also disrupted head–body synchronization and led to prolonged head muscle activation during both crawling and swimming. Importantly, SMB ablation increased global muscle Ca²⁺ activity, indicating that SMB neurons normally act to prevent excessive or sustained muscle activation. These findings suggest that the SMB neurons do not simply drive locomotion, but regulate curvature switching by limiting muscle activation amplitude and duration. In this model, the SMB neurons function as a brake-like modulatory node that stabilizes posture transitions during swimming. By suppressing persistent head muscle activity, the SMB neurons enable rapid curvature reversal and maintain oscillatory symmetry. This role is conceptually distinct from the SMD neurons, which primarily influence curvature switching kinetics without affecting overall muscle activation levels. Thus, the SMB neurons appear to couple head posture control with global motor state transitions, serving as a critical coordinator that aligns head output with whole-body oscillatory demands during swimming. This organizational logic parallels vertebrate locomotor control, in which brainstem locomotor regions such as the cuneiform nucleus and pedunculopontine nucleus modulate gait transitions and stabilize locomotor states through distributed excitatory networks ^8,10^. In this context, the SMB neurons may represent a simplified gating module that constrains motor output to enable state-specific oscillatory patterns, reflecting a conserved principle in which defined neural nodes coordinate transitions between locomotor patterns.

NALCN channel complexes have been broadly implicated in sustaining neuronal excitability and rhythmic motor output across species. In *C. elegans*, the NCA channel complex maintains persistent motor circuit activity ^49,50^. Our findings refine this framework by showing that UNC-79–dependent NALCN signaling is required to stabilize the swimming motor state. Although *unc-79* mutants retained the ability to generate individual body bends, they exhibited pronounced delays in gait switching and reduced swimming frequency, indicating that NALCN activity does not encode bend generation itself but instead supports the excitability necessary to sustain a high-frequency locomotor state. The requirement for UNC-79 across multiple head neuronal hubs, including the RMD motor neurons and the RID interneuron, further suggests that motor state stabilization emerges from coordinated modulation of distributed circuit excitability rather than from a single command neuron. This circuit-level stabilization must ultimately be translated into the muscle activation patterns that execute the swimming motor state. Our data position UNC-29, a muscle-expressed nicotinic acetylcholine receptor subunit, as a critical effector of this transformation. *unc-29* mutants exhibited delayed gait transitions and defective swimming accompanied by elevated mean body-wall Ca^2+^ levels despite slowed propagation, indicating a mismatch between synaptic drive and organized muscle calcium dynamics. These observations suggest that UNC-29 contributes to calibrating how excitatory input is converted into state-appropriate muscle activation. The paradoxical increase in Ca^2+^ may arise from prolonged depolarization or compensatory recruitment of parallel excitatory pathways that are normally balanced by nAChR-mediated currents. Differential phenotypes among L-AChR subunit mutants further imply that UNC-29 may participate in receptor assemblies optimized for the demands of the swimming motor state. Together, these findings support a hierarchical model in which UNC-79 stabilizes circuit excitability, while UNC-29 tunes muscle responsiveness to ensure coherent execution of the swimming motor state.

Our findings converge on a hierarchical model in which gait transition emerges from coordinated interactions between head motor neurons, interneuron excitability modules, and muscle-intrinsic receptor dynamics (Figure 7F). First, the SMB head motor neurons function as a gating node that suppresses persistent head muscle activation and aligns head output with whole-body oscillatory patterns during swimming. Second, UNC-79–dependent NALCN signaling operates across distributed interneuron hubs to stabilize the high-excitability state required for sustained swimming. Third, UNC-29–containing nAChRs in body wall muscles refine calcium amplitude and propagation dynamics to ensure efficient curvature switching. Rather than being driven by a single trigger neuron or molecular switch, gait transition appears to represent a systems-level state reconfiguration involving coordinated modulation of excitability, temporal precision, and muscle gain control. This distributed architecture may confer robustness, allowing animals to flexibly adapt motor patterns across environmental contexts. Conceptually, our study links defined head motor neurons to downstream molecular effectors and muscle calcium dynamics, providing a mechanistic bridge between neural circuit activity and biomechanical output. Given the evolutionary conservation of NALCN channels and nicotinic acetylcholine receptors, similar circuit-to-muscle signaling principles may operate in more complex locomotor systems, where transitions between motor states require synchronized modulation of excitability and muscle activation patterns.

## STAR★METHODS

### KEY RESOURCES TABLE

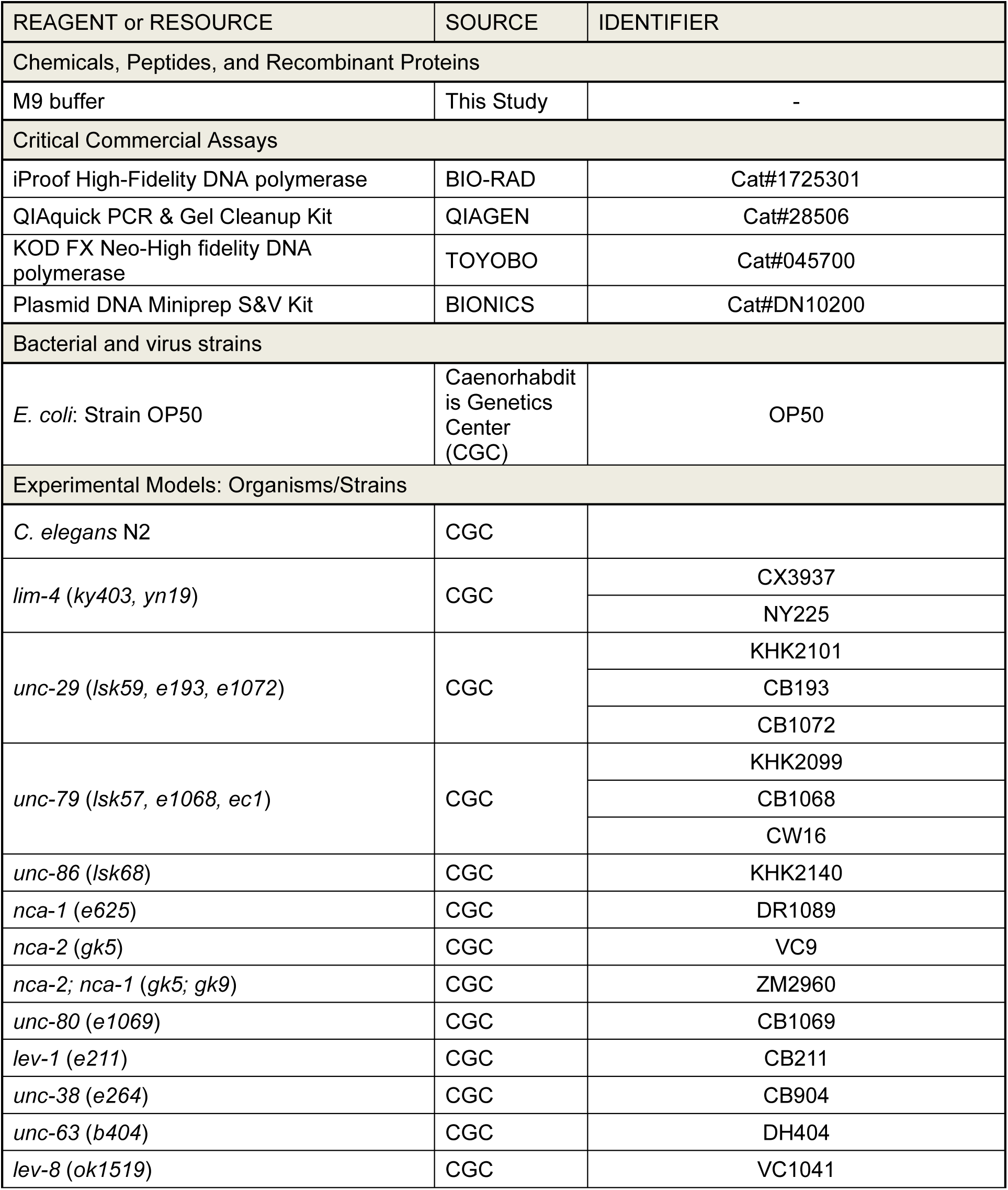
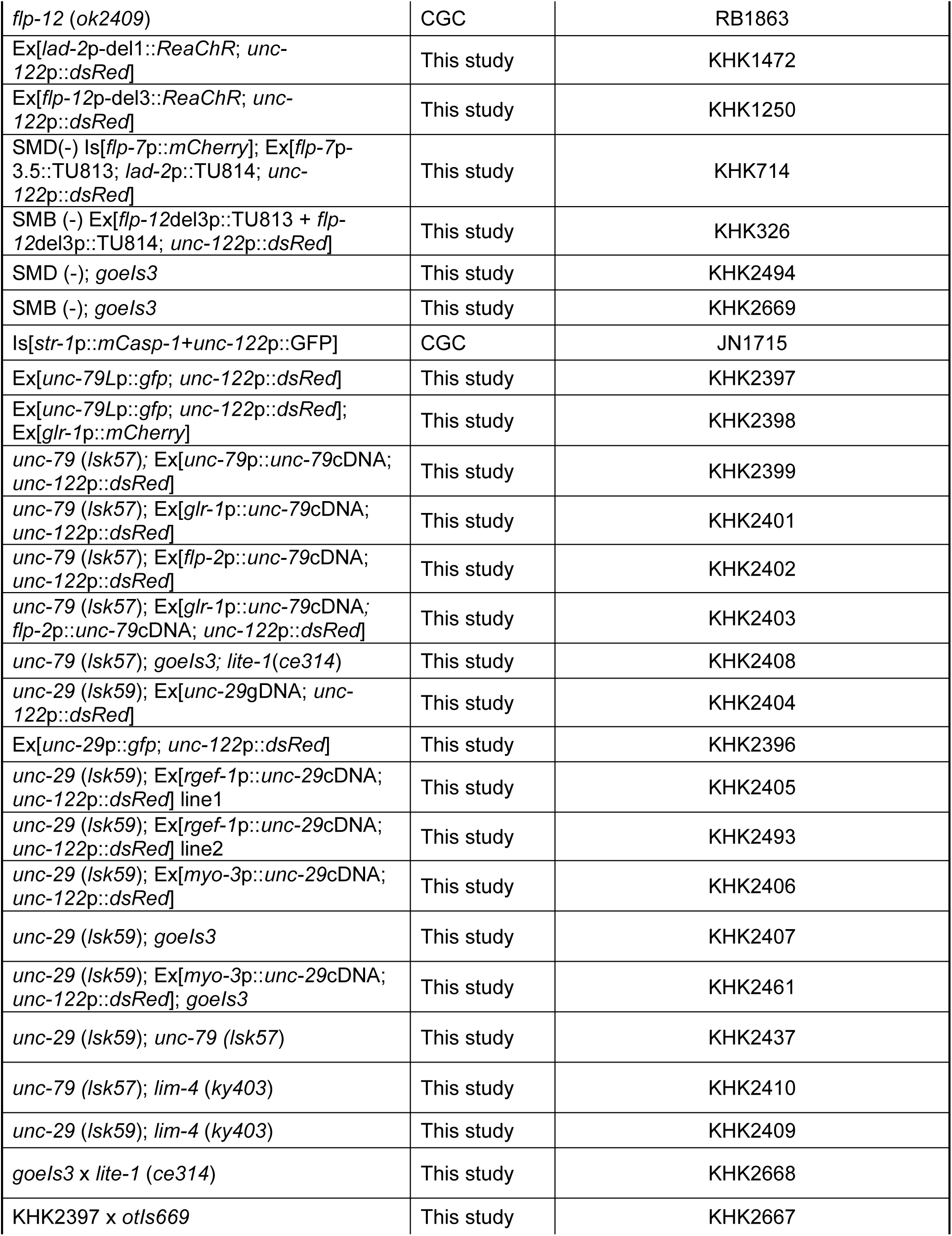
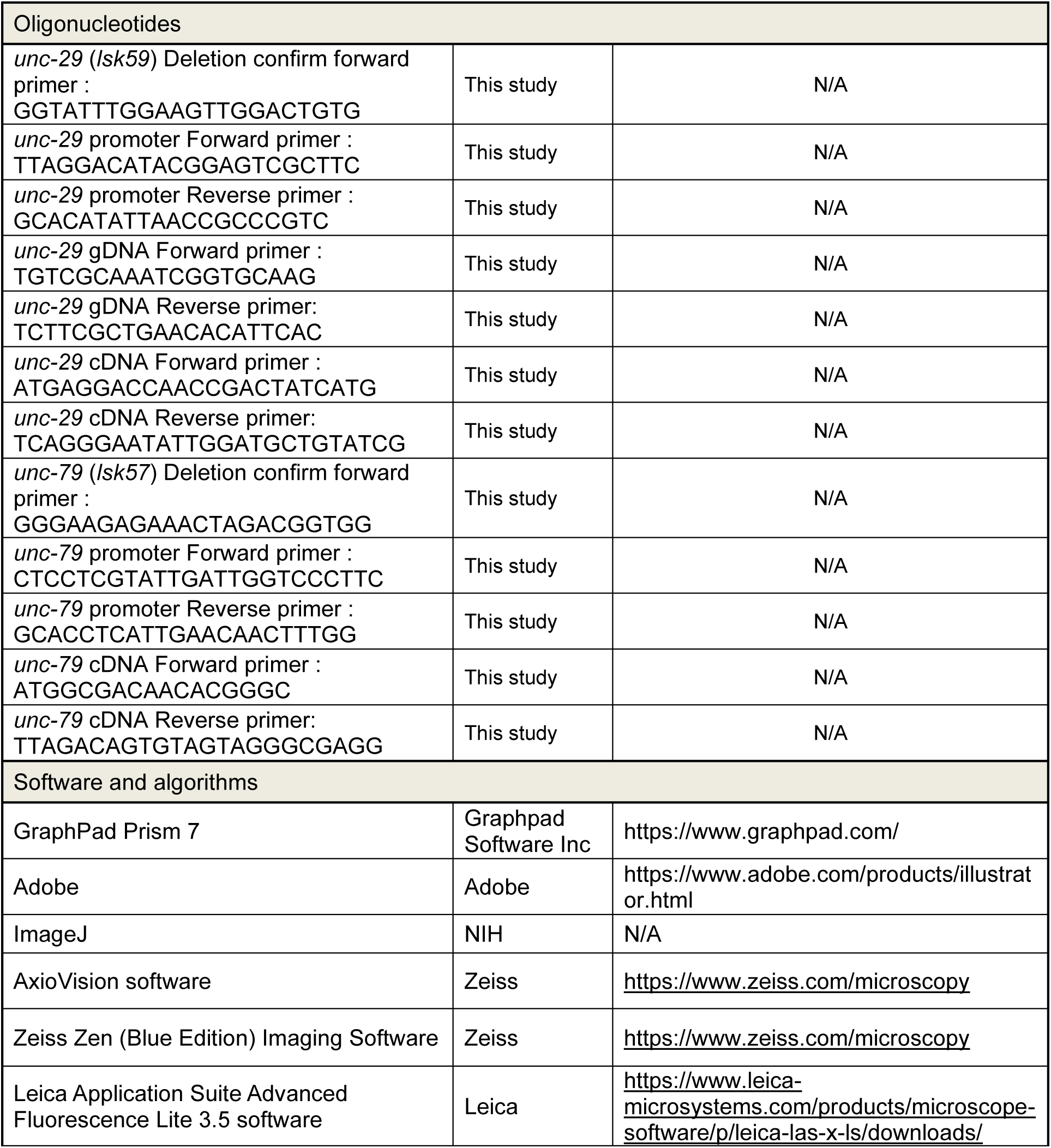

### RESOURCE AVAILABILITY

#### Lead contact

Further information and requests for resources and reagents should be directed to and will be fulfilled by the Lead Contact, Kyuhyung Kim.

#### Materials availability

Materials generated in this study, including strains, plasmids, and clones, are freely available from the Lead Contact upon request.

#### Data and code availability

This study did not generate any unique datasets or code.

### METHOD DETAILS

#### Strains

All strains were maintained at 20℃ ^51^. The N2 Bristol was used as a wild-type strain. Mutant strains and transgenic strains used in this study are listed in key resource table.

#### Isolation of gait transition mutants

The N2 Bristol was used to perform EMS mutagenesis according to Brenner ^20^. Three alleles (*lsk57*, *lsk59* and *lsk67*) in which exhibited gait transition defects were isolated and were genetically mapped on LGIII, LGI, and LGIII, respectively. The molecular lesions of *lsk57*, *lsk59* and *lsk67* were further identified by whole-genome sequencing (BGI, see below).

#### Whole genome resequencing of mutant allele and data analysis

Adult wild-type and gait transition mutant animals (*lsk57*, *lsk59* and *lsk67*) were cultured in NGM plates with OP50 at 20℃ for 4-5days. After harvest, worms were lysed and treated with Puregene Tissue Kit (Qiagen) to extract the whole genomic DNA ^52^. Purified genomic DNA of the samples was re-sequenced at the Beijing Genomics institute. The coverage of whole genome re-sequencing was 50Χ and the raw sequencing data were preprocessed and aligned onto *C. elegans* reference genome (WBcel235/ce11 assembly) using IGV (Integrative Genomics Viewer) ^53^. Genomic variants were annotated using SNP data, and candidate mutations were filtered based on the presence of premature translation stop codons. Through this analysis, *unc-79* was identified in *lsk57*, *unc-29* in *lsk59*, and *unc-86* in *lsk67*.

#### Molecular biology and transgenic worms

The pPD95.77 vector and its derivatives were used for the worm constructs in this study. For promoter fusions, *unc-79* (3.6kb) and *unc-29* (3.4kb) promoter regions were amplified by PCR from N2 genomic cDNA. Green fluorescent protein (GFP) fused with *unc-79* and *unc-29* promoter by using various enzymes. To generate transgenic worms, 50ng of each reporter construct was injected, with 50ng of *unc-122*p::*dsRed* as an injection marker. For the *unc-79* and *unc-29* rescue experiments, the *unc-79*, *glr-1* (5.8kb) and *flp-2* (2.3kb) promoters were fused with *unc-79* cDNA and the *unc-29*, *myo-3* (2.3kb) and *rgef-1* (583bp) promoters were fused with *unc-29* cDNA isolated from N2 cDNA library. To generate rescue lines, 75ng *unc-79* rescue constructs injected into *unc-79* (*lsk57*) and 50ng *unc-29* rescue constructs injected into *unc-29* (*lsk59*) with 50ng of *unc-122*p::*dsRed* as an injection marker.

#### Behavior recording and analysis

To record animal locomotion, a well-fed young adult worm was transferred to an empty NGM plate and allowed to acclimate for 1 min. Movies were recorded at 25 frames/s using a Nikon digital microscope equipped with HDMI camera. Gait transitions were recorded after the 1-min acclimation period on an empty NGM plate. Subsequently, 1 μL of M9 buffer was dropped beside the worm’s body, allowing the worm to move into the liquid puddle. Video recording continued for 3 min after the worm approached the puddle. Recorded videos were analyzed using ImageJ software and the Tierpsy Worm Tracker ^54^. Seven behavioral phenotypes were analyzed: crawl-to-swim onset, crawling frequency, swimming frequency, curvature switch time, posture holding time, arc length change, and head–vulva–tail angle. Detailed analysis methods are described below.

Crawl-to-swim transition time (C–S transition time) was defined as the time from initial contact with the liquid to the onset of swimming behavior. From sequential images, the transition time was calculated as: (end frame-start frame) x (1/25). The start frame was defined as the frame immediately before the worm approached the liquid, and the end frame was defined as the frame at which the worm initiated swimming with a characteristic C-shaped posture and forward movement.

Crawling frequency and swimming frequency were defined as the number of sinusoidal crawling bends or C-shaped swimming bends per second, respectively. From recorded movies, the total number of crawling or swimming bends was counted and divided by 30 s.

Curvature switch time was defined as the time required for the worm to transition from one C-shaped body bend to the opposite curvature. Curvature switch time was calculated using the following formula: (stop opposite-side bending frame – start bending frame) x (1/25).

Posture holding time was defined as the duration for which the worm maintained a fully bent C-shaped posture before transitioning to the opposite bend. Posture holding time was calculated as: (approach-to-bend frame – start opposite-side bending frame) x (1/25).

Arc length change was defined as the change in body arc length during crawling and swimming. Differential arc length was calculated using the following formula: (average minimum length)/ (average maximum length). Raw arc length data (annotated as “*body length*” in Tierpsy) were obtained by processing videos of crawling and swimming behavior using the open-source Tierpsy platform.

The head–vulva–tail (three-point) angle was defined as the average angle formed by the head, vulva, and tail during one cycle of C-shaped bending. Angles were measured using the angle tool in ImageJ software, and angle values were averaged across nine consecutive frames corresponding to a single bending cycle (0–2π interval).

#### Calculation of Pearson’s correlation coefficient

The Pearson correlation coefficient (r) was used to quantify the linear relationship between two variables. For two datasets X=(x1,x2,…,xn) and Y=(y1,y2,…,yn), the Pearson correlation coefficient was calculated as:

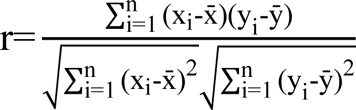

where x and y represent the mean values of X and Y respectively. The coefficient 𝑟ranges from −1 to +1, with values close to +1 or −1 indicating strong positive or negative linear correlations, respectively, and values close to 0 indicating little or no linear correlation. Pearson correlation analysis was performed using Microsoft excel worksheet.

#### Calcium imaging and image analysis

*In vivo* calcium imaging of body wall muscles was performed by measuring changes in fluorescence of the calcium indicator GCaMP3.35, as previously described ^48^. Well-fed young adult–stage worms were used to observe body wall muscle calcium activity. For freely moving conditions, each worm was transferred to an empty NGM plate and recorded after a 1-min acclimation period. Crawling behavior was filmed for 30 s, after which M9 buffer was dropped beside the worm’s body to induce gait transition and swimming behavior. The *C. elegans* transgenic strains used for calcium imaging experiments included HBR4, KHK2407, KHK2461, and KHK2408. All videos were recorded at 20 frames/s using a Leica high-performance fluorescence stereomicroscope (M205FA) with Leica Application Suite Advanced Fluorescence Lite 3.5 software and were analyzed using ImageJ software. To quantify calcium signals from sequential images, segmented lines were drawn over the regions of interest (ROIs) corresponding to muscles in each frame, and the average pixel intensity was measured. In Figures 1E, 3A, 3C, S2B, S2C, 5G, and 7A–B, ROIs corresponded to ventral body wall muscles, whereas in Figures 3B and 3D, ROIs corresponded to head muscles.

#### Optogenetics

L4 transgenic worm larvae expression ReaChR::mKate2 trangenes under the control of indicated promoters *flp-12*del3p (KHK1250) and *lad-2*del1p (KHH1472)) were transferred 12h before the assay to either normal OP50 plates or OP50-retinal plates containing 1mM all-trans-retinal (ATR, Sigma). OP50-retinal plates were prepared by seeding 200μL op50 with 2μL 100mM ATR (Sigma) in 100% ethanol. To stimulate the ReaChR, 617nm LED was illuminated using Fiber-Coupled LED (M565F3, THORLABS) at roughly 0.05mW/mm as measured with an optical power/energy meter. Transferred worms were recorded in at least 3min, and red light was exposed during 2min before/after M9 buffer dropping. After recording, crawling and swimming frequency were analyzed by image-J software.

#### Confocal imaging and Cell ID

To identify neuronal sites of gene expression, we used young-adult hermaphrodites. We placed three worms on a 3% agar pad on the cover glass and paralyzed the worms using 1M sodium azide (Sigma-Aldrich). All imaging was performed with an LSM780 confocal micro-scope (Carl Zeiss) at room temperature (22-25℃) ^55^. NeuroPAL cell identification was done by assessing position and size using Nomarski optics and by crossing with the NeuroPAL strain (*otIs669*) ^39^. Imaging experiments using the NeuroPAL line were conducted with specific laser settings, which are available for download on yeminilab.com. Analysis of NeuroPAL images for cell identification was conducted as in Yemini et al. (2021).

#### Statistical analysis

Plots were generated in GraphPad Prism software. Comparisons and P-value calculations were made between animals of the same or different strains, and treated and untreated animals, using Student’s t-test, one-way ANOVA and two-way ANOVA with corrections for multiple comparison testing. More statistical information is represented in all figure legends.

## Acknowledgements

We are grateful to the Caenorhabditis Genetics Center (NIH Office of Research Infrastructure Programs, P40 OD010440) and the National BioResource Project (Japan) for strains. We are also grateful to KyeongJin Kang, Hyun-Ho Lim, and K. Kim Labs for helpful discussion and/or critical comments on this manuscript. This work was supported by the National Research Foundation of Korea (RS-2024-00356180 and RS-2024-00467236), the DGIST International Joint Research Project (24-KUJoint-07), and the KBRI (25-BR-05-07) to K. Kim.

## AUTHOR CONTRIBUTIONS

K.M. M. conceived and designed the study. K.M. M. performed the experiments. J. K and J. C assisted with EMS mutagenesis screening. K.M. M and K.K. analyzed the data. K.M. M., and K.K. wrote the paper. All authors read, edited and approved the final manuscript.

