## Supplementary Figures for "A circuit-to-muscle signaling axis controls locomotor gait transitions in *C. elegans*"

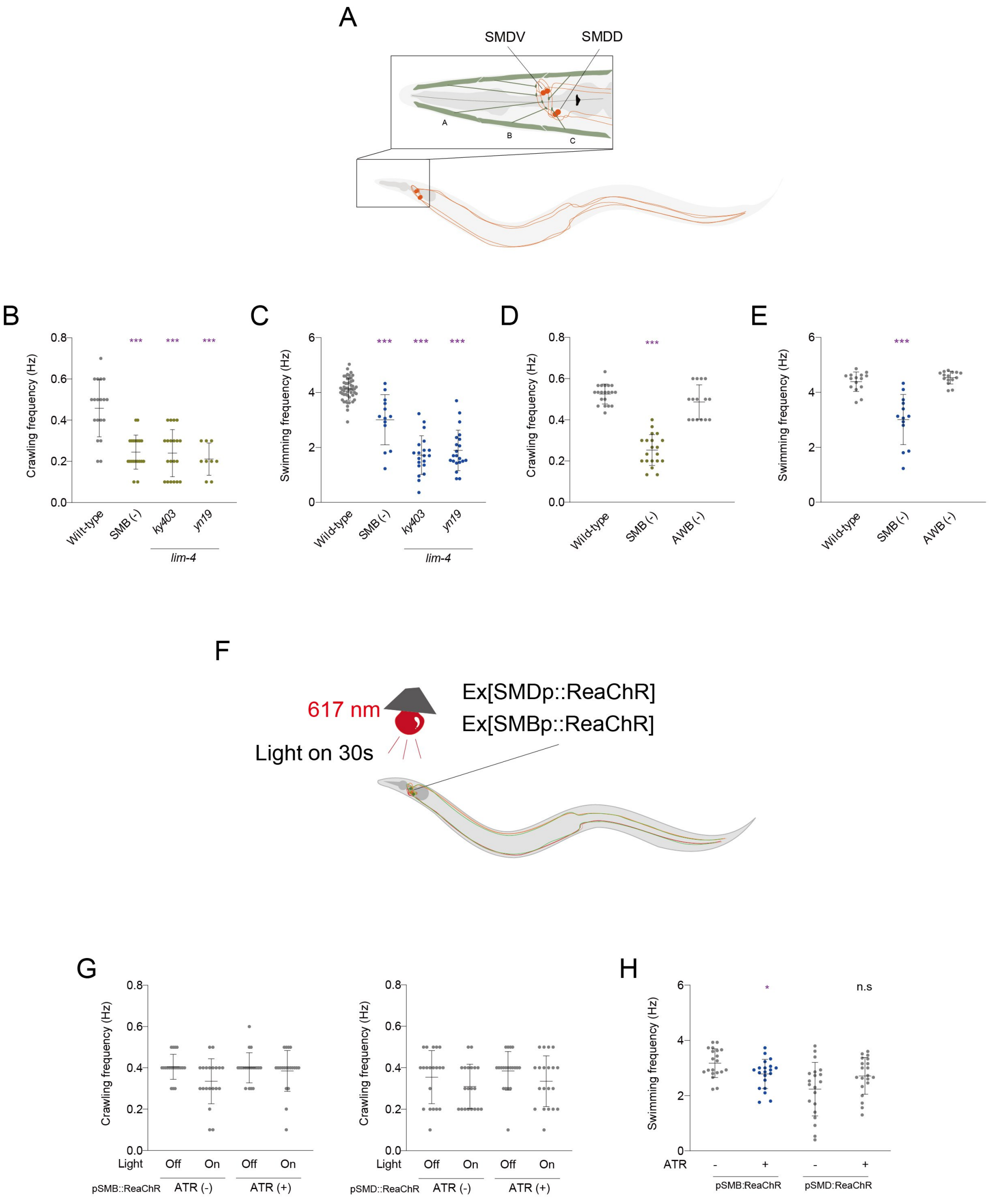


**Figure S1. Head motor neurons SMB and SMD differentially regulate swimming behavior Related to Figure 2.**
(A) Schematic illustrating the SMD head motor neurons in *C. elegans*. (B, C) C-S transition time and swimming frequency of SMB defective *lim-4* mutants in two alleles (*ky403* and *yn19*). n ≥ 9 for each. (D, E) C-S transition time and swimming frequency of wild-type, SMB (-) (KHK340) and AWB (-) (JN1715) worms. n ≥ 13 for each. (F) Schematic drawing of optogenetic experiments targeting the SMB and the SMD neurons. (G, H) C-S transition time and swimming frequency of SMB, SMD-specific optogenetic animals under red light (617 nm) stimulus. SMB-, SMD-specific optogenetic animals expressing red-shifted channelrhodopsin (ReaChR) under the control of the *lim-4* and *lad-2* gene promoters, respectively. n ≥ 20 for each. Data are represented ± SD. *, *** indicate significant differences from wild-type at *$p < 0.05$ and ***$p < 0.001$ are shown (one-way ANOVA test followed by Dunnett's multiple comparisons test).

Figure S2 Moon et al.

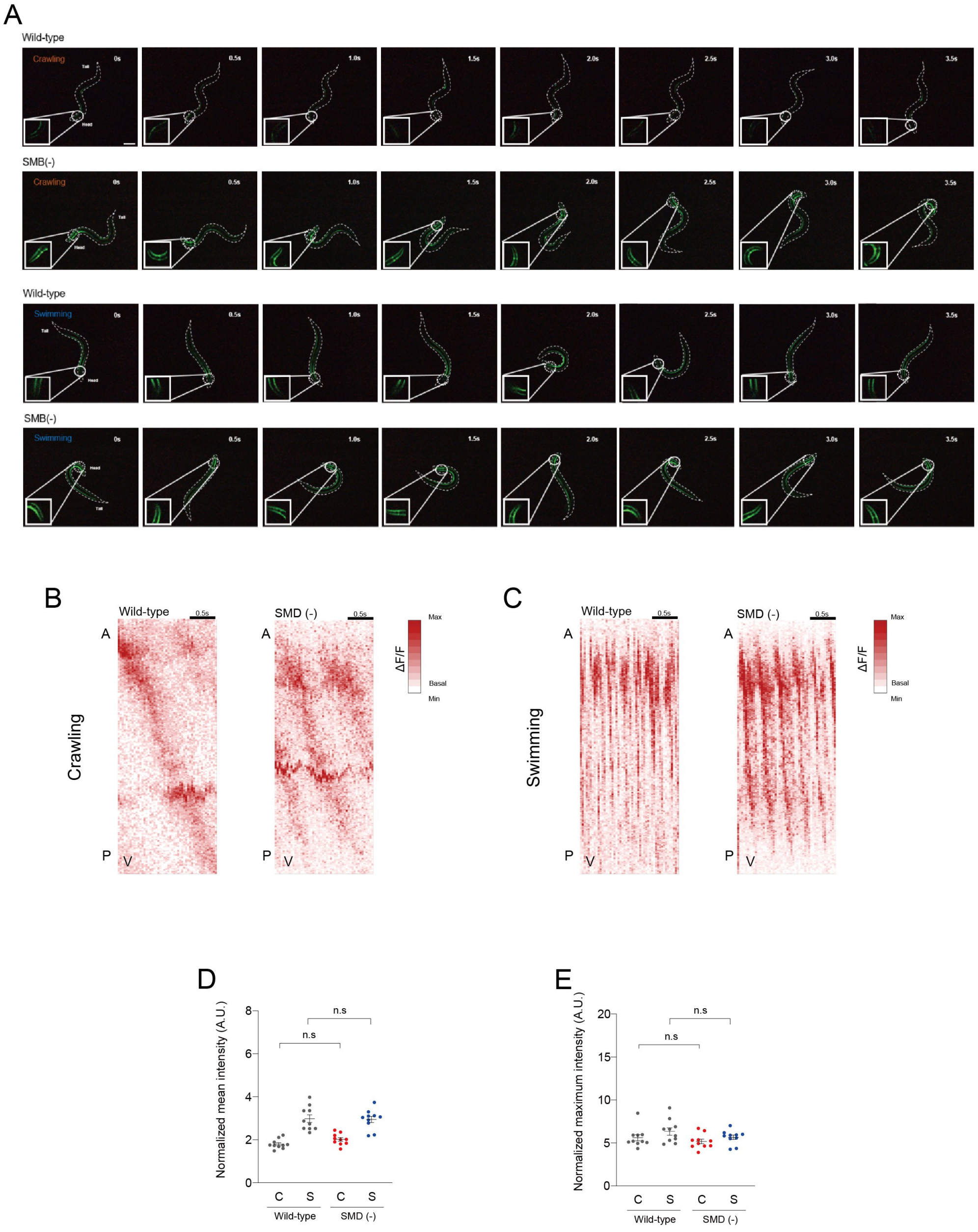


**Figure S2. The SMB neurons regulate muscle contraction and locomotion state transition Related to Figure 3.**

(A) Representative time-series images of adult animals expressing *myo-3*p::GCaMP3.35 in wild-type and SMB (-) worms during crawling and swimming. Scale bar = 100 μm (B, C) Representative $Ca^{2+}$ activity maps of ventral body-wall muscles in wild-type and SMD (-) (KHK1174) animals during crawling and swimming, respectively. The x-axis denotes time, and the y-axis indicates body position from anterior to posterior. (D, E) Mean and maximum $Ca^{2+}$ fluorescence intensity of body-wall muscles in wild-type and SMD (-) animals during crawling and swimming. Data are represented ± SEM. n.s indicate non-significant difference from wild-type is shown (one-way ANOVA test followed by Tukey's multiple comparisons test).

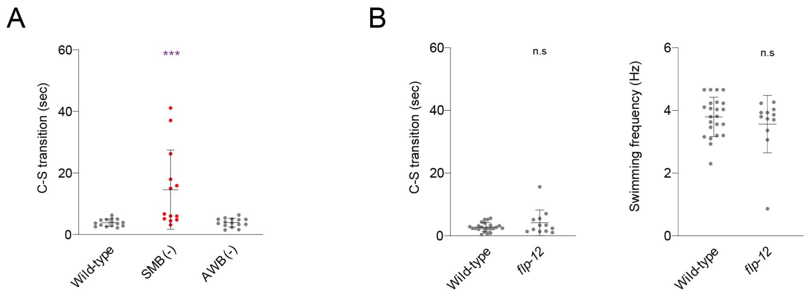


**Figure S3. The SMB neurons regulate muscle contraction and locomotion state transition Related to Figure 3.**
(A) C-S transition time of SMB (-) and AWB (-). n ≥ 13 for each. (B) C-S transition time and swimming frequency of wild-type and *flp-12* mutants. n ≥ 12 for each. Data are represented ± SD. *** indicates a significant difference from wild-type at ***$p < 0.001$ (one-way ANOVA test followed by Dunnett's multiple comparisons test).

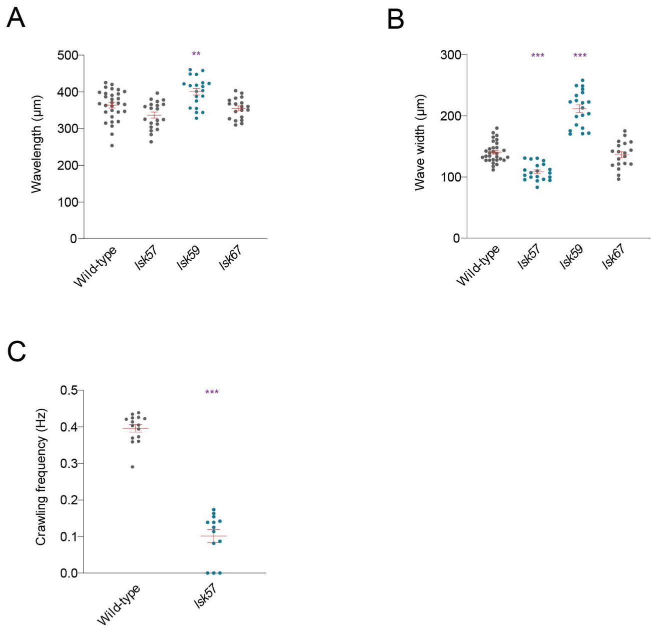


**Figure S4. Isolation of mutant strains exhibiting abnormal gait transitions**
**Related to Figure 4.**
(A, B) Mean wave width and wavelength of wild-type, *lsk57*, *lsk59,* and *lsk67* animals. n ≥ 8 for each. (C) Crawling frequency of wild-type and *lsk57* animals after multiple generations. n ≥ 13 for each. Data are represented ± SD (C, D) and SEM (A, B). ** and *** indicate a significant difference from wild-type at **$p < 0.01$ and ***$p < 0.001$ are shown (one-way ANOVA test followed by Dunnett's multiple comparisons test).

Figure S5

Moon et al.

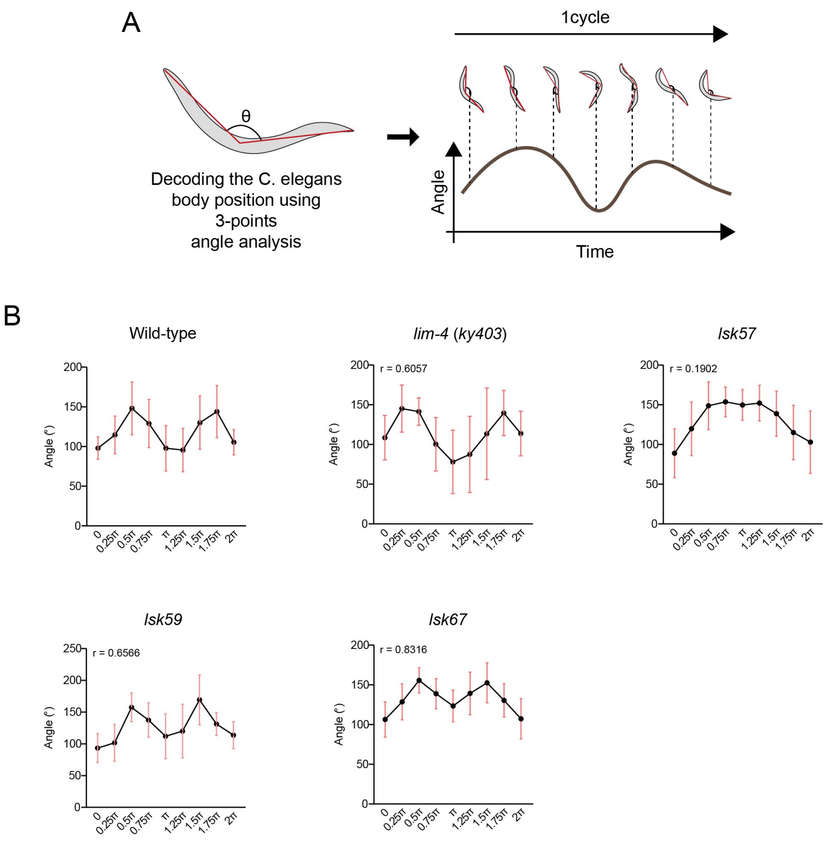


**Figure S5. Isolation of mutant strains exhibiting abnormal swimming posture**
**Related to Figure 4.**

(A) Schematic illustrating the head–vulva–tail (three-point) angle analysis used to quantify swimming posture (see Methods). (B) Three-point angle analysis of wild-type animals and isolated mutant strains during swimming. The x-axis represents one swimming cycle, and the y-axis indicates the angle derived from the three-point analysis. n =10 for each. Pearson correlation coefficients (r) were calculated to assess waveform similarity (see Methods).

Figure S6

Moon et al.

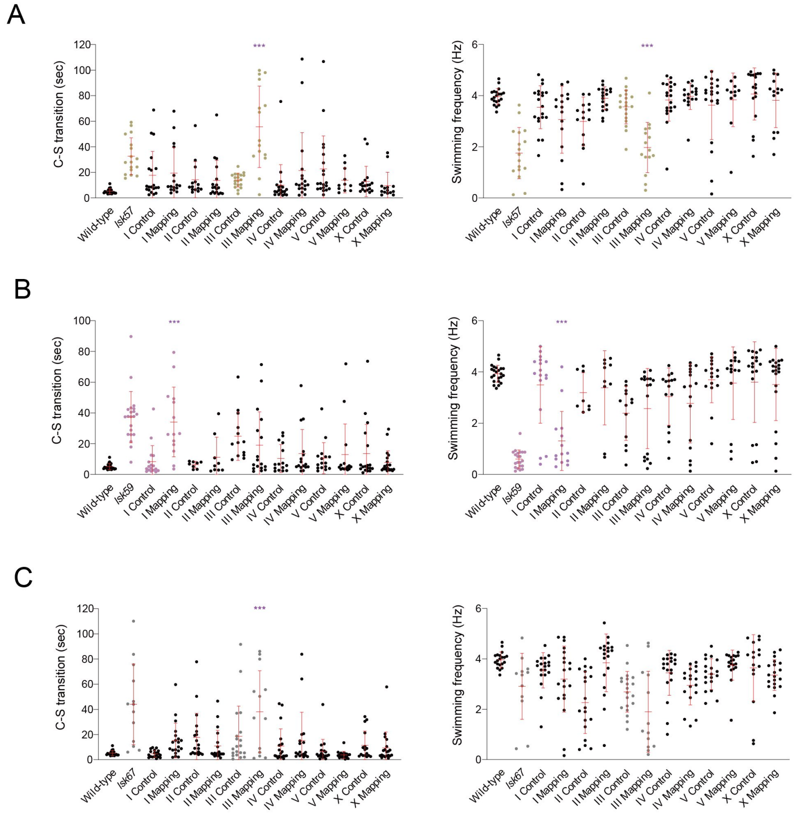


**Figure S6. Chromosomal mapping results for isolated mutant strains**
**Related to Figure 4.**

(A-C) Crawl-to-swim transition time and swimming frequency of isolated mutant strains and mutants crossed with chromosomal marker containing transgenic animals. n ≥ 13 for each. Data are represented ± SD. *** indicates a significant difference from wild-type at ***$p < 0.001$ (one-way ANOVA test followed by Dunnett's multiple comparisons test).

Figure S7



A

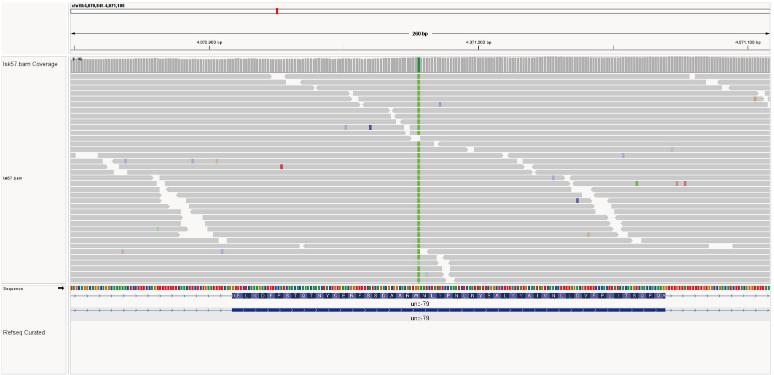


B

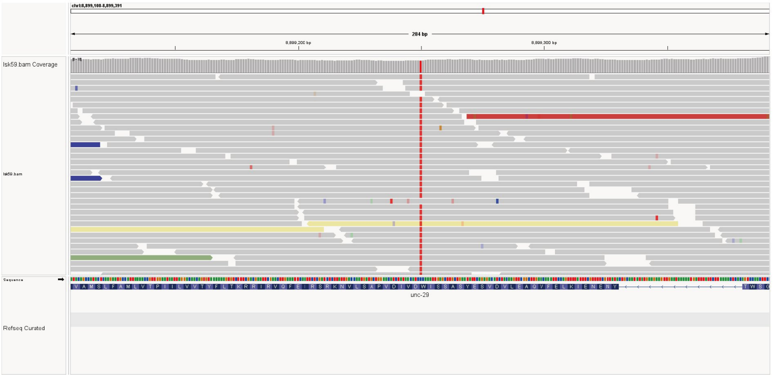


C

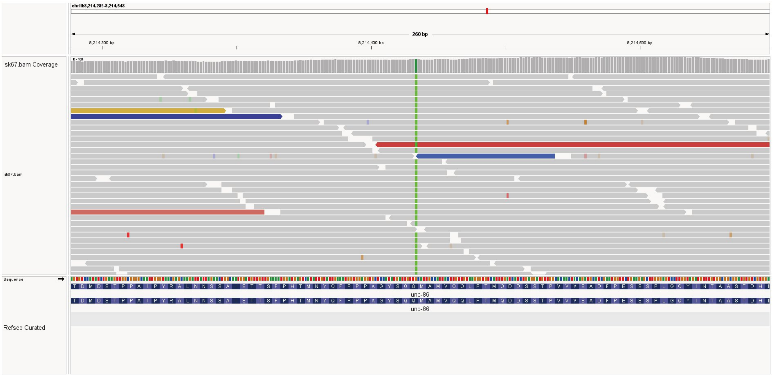


**Figure S7. Whole genome sequencing results for isolated mutant strains**
**Related to Figure 4.**
(A–C) Stop-gain mutation sites identified in isolated mutant strains. Whole-genome sequencing data are shown as Integrative Genomics Viewer (IGV) tracks.

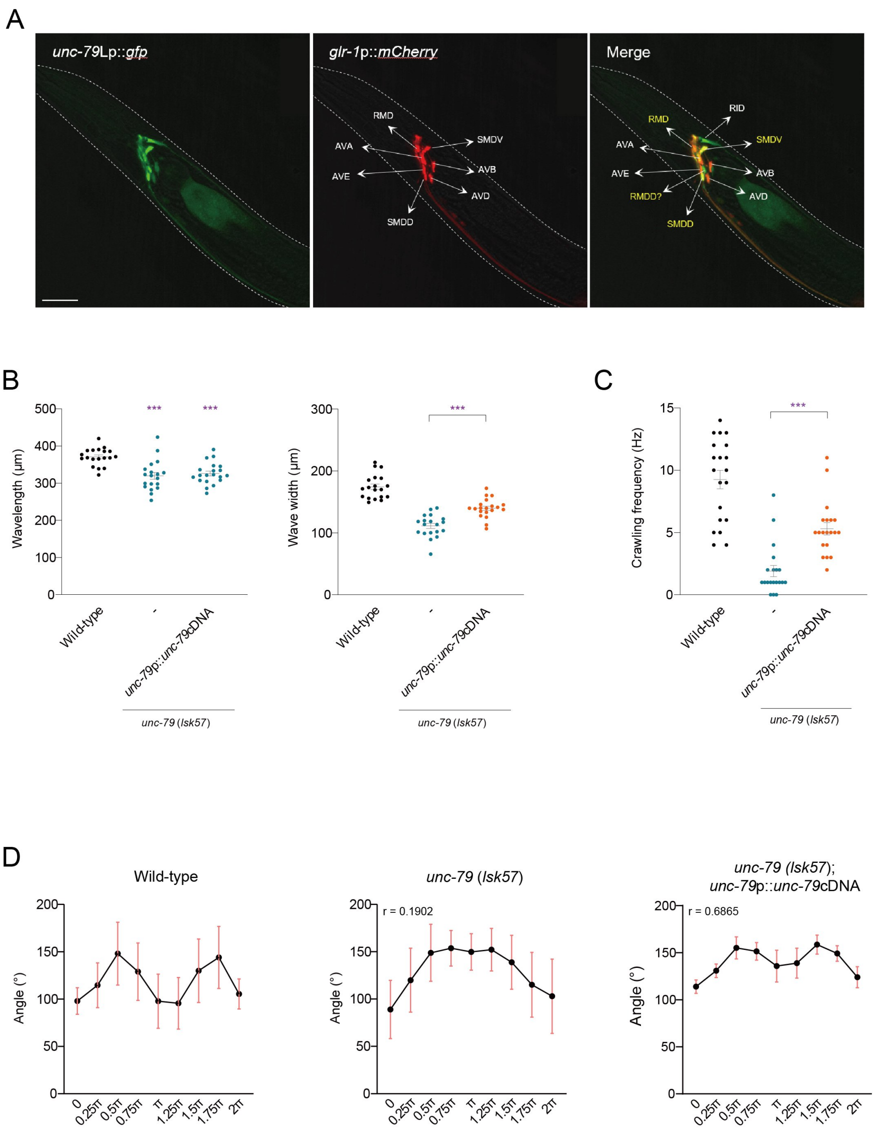


**Figure S8. UNC-79-expressing interneurons mediate gait transition and swimming behavior Related to Figure 5.**

(A) Representative images of a wild-type animal co-expressing *unc-79*p::*gfp* and *glr-1*p::*mCherry* (left : *unc-79*p::*gfp*, middle : *glr-1*p::*mCherry*, right: merged). The *unc-79* promoter drives expression in RMD, RID, and SMDD neurons. Anterior is to the left. Scale bar = 20μm. (B) Mean wave width and wavelength of wild-type and *lsk57* mutant animals expressing *unc-79* cDNA under the control of the *unc-79* long-isoform promoter. n ≥ 19 for each. (C) Crawling frequency of wild-type and *lsk57* mutant animals expressing *unc-79* cDNA under the control of the *unc-79* long-isoform promoter. n = 20 for each. (D) Three-point angle analysis of wild-type and *lsk57* mutant animals expressing *unc-79* cDNA under the control of the *unc-79* long-isoform promoter during swimming. n = 10 for each. Data are represented ± SEM. *** indicates a significant difference from wild-type at ***$p < 0.001$ (one-way ANOVA test followed by Tukey's multiple comparisons test).

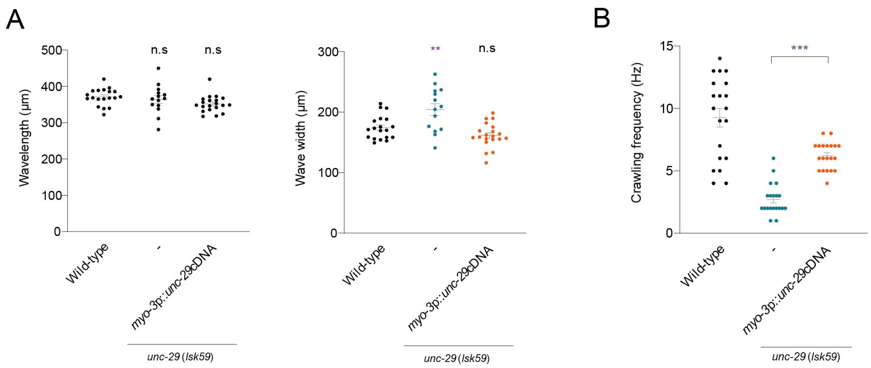


**Figure S9. *unc-29* cDNA expression in body wall muscle restores crawling defects**

**Related to Figure 6.**

(A) Mean wave width and wavelength of wild-type and *unc-29* mutant animals expressing *unc-29* cDNA under the control of the *myo-3* promoter. n ≥ 14 for each. (B) Crawling frequency of wild-type and *unc-29* mutant animals expressing *unc-29* cDNA under the control of the *myo-3* promoter. n = 20 for each. Data are represented ± SEM. ** and *** indicate a significant difference from wild-type at **p < 0.01 and ***p < 0.001 (one-way ANOVA test followed by Tukey's multiple comparisons test).

**Supplementary Video 1, 2 and 3**
**Related to Figure 2.**
Swimming behavior of wild-type (Supplementary Video 1), SMB (-) (Supplementary Video 2) and SMD (-) (Supplementary Video 3) animals. Animals were recorded for 10 seconds in M9 buffer.
